# Modulation of flight and feeding behaviours requires presynaptic IP_3_Rs in dopaminergic neurons

**DOI:** 10.1101/2020.08.19.258400

**Authors:** Anamika Sharma, Gaiti Hasan

## Abstract

Innate behaviours, though robust and hard wired, rely on modulation of neuronal circuits, for eliciting an appropriate response according to internal states and external cues. *Drosophila* flight is one such innate behaviour that is modulated by intracellular calcium release through inositol 1,4,5-trisphosphate receptors (IP_3_Rs). Cellular mechanism(s) by which IP_3_Rs modulate neuronal function for specific behaviours remain speculative, in vertebrates and invertebrates. To address this, we generated an inducible dominant negative form of the IP_3_R (IP_3_R^DN^). Flies with neuronal expression of IP_3_R^DN^ exhibit flight deficits. Spatiotemporal expression of IP_3_R^DN^ helped identify key flight-modulating dopaminergic neurons with axonal projections in the mushroom body. Attenuation of IP_3_R function in these presynaptic dopaminergic neurons resulted in flies with shortened flight bouts and a disinterest in seeking food, accompanied by reduced excitability and dopamine release upon cholinergic stimulation. Our findings suggest that the same neural circuit modulates the drive for food search and for undertaking longer flight bouts.

## Introduction

The Inositol-1, 4, 5-trisphosphate receptor (IP_3_R) is an Endoplasmic Reticulum (ER) resident ligand-gated calcium (Ca^2+^) channel found in metazoans. In neurons, the IP_3_R is activated through multiple classes of signaling molecules that include neuromodulators such as neuropeptides, neurotransmitters and neurohormones. Studies from ex-vivo vertebrate neurons have identified a role for the IP_3_R in multiple cellular processes including regulation of neurite growth (Takei et al., 1998; Xiang et al., 2002), synaptic plasticity (Fujii et al., 2000; Nishiyama et al., 2000) and more recently pre-synaptic neurotransmitter release (Gomez et al., 2020). However, the relevance of IP_3_-mediated signaling mechanisms to cellular processes and subsequent behavioural and neurophysiological outputs need better understanding. In non-excitable cells Ca^2+^ release through the IP_3_R regulates a range of cellular events including growth, secretion (Inaba et al., 2014), gene expression (Ouyang et al., 2014) and mitochondrial function (Cárdenas et al., 2010, Bartok et al., 2019). In excitable cells such as neurons, identifying cellular functions of the IP_3_R is more complex because in addition to IP_3_-mediated Ca^2+^ release, there exist several plasma-membrane localised ion channels that bring in extracellular Ca^2+^ in response to neurotransmitters and changes in membrane excitability.

Three IP_3_R isotypes, IP_3_R1, 2 and 3 are encoded by mammalian genomes (Furuichi et al., 1994; Taylor et al., 1999). Of these, IP_3_R1 is the most prevalent isoform in neurons and is relevant in the context of several neurodegenerative disorders (Terry et al., 2019). Human mutations in IP_3_R1 cause Spinocerebellar ataxia 15 (SCA15), SCA29 and Gillespie syndrome (Hasan & Sharma, 2020). Disease causing IP_3_R1 mutations span different domains but several are clustered in the amino terminal IP_3_ binding region, from where they impact ER-Ca^2+^ release when tested in mammalian cell lines (Ando et al., 2018). A common feature of these neurological disorders is loss of motor-coordination or ataxia. The ataxic symptoms arise primarily from malfunction and/or degeneration of cerebellar Purkinje neurons where the IP_3_R1 is expressed abundantly in the soma, dendrites and axons. Dendritic expression of the IP_3_R in Purkinje neurons determines Long Term Depression (LTD), a form of post-synaptic plasticity (Miyata et al., 2000). Interestingly, a recent study in *Drosophila* also identified the IP_3_R as an essential component of post-synaptic plasticity, required for decoding the temporal order of a sensory cue and a reward stimulus, in neurons of a higher brain centre, the Mushroom Body (Handler et al., 2019). Somatic expression of the IP_3_R very likely contributes to maintenance of cellular Ca^2+^ homeostasis (Berridge, 2016). A pre-synaptic role for the IP_3_R has been demonstrated in the *Drosophila* neuromuscular junction (Shakiryanova et al., 2011), motor neurons (Klose et al., 2010) and most recently in axonal projections of Purkinje neurons (Gomez et al., 2020). Physiological and behavioural significance of pre-synaptic IP_3_/Ca^2+^ signals in either case however remain speculative.

To understand how IP_3_R alters neuronal function and related neurophysiology and behaviour we have in the past studied several mutants for the single IP_3_R gene (*itpr*) in *Drosophila melanogaster* (Joshi et al., 2004). The *Drosophila* IP_3_R shares 60% sequence identity, similar domain organisation, biophysical and functional properties with mammalian IP_3_Rs(Chakraborty & Hasan, 2012; Srikanth & Hasan, 2004). Several hypomorphic and heteroallelic combinations of *Drosophila* IP_3_R mutants are viable and exhibit flight deficits ranging from mild to strong, the focus of which lies in aminergic neurons (Banerjee et al., 2004). However, such mutants do not easily allow cell specific attenuation of IP_3_R function. In this context, a dominant-negative mutant form of mammalian IP_3_R1, generated recently, abrogated IP_3_R function in mammalian cell lines (Alzayady et al., 2016). The mammalian dominant negative IP_3_R1 gene was based on the finding that all monomers in the IP_3_R tetramer need to bind IP_3_ for channel opening (Alzayady et al., 2016). Failure of a single IP_3_R monomer to bind IP_3_ renders the resultant IP_3_R tetramer non-functional. *Drosophila* IP_3_Rs are also tetrameric suggesting that a similar strategy for generating a dominant-negative IP_3_R could be used for cell-specific studies of neuronal function and behaviour. Consequently we generated and characterised a *Drosophila* IP_3_R dominant negative (IP_3_R^DN^) transgene. Cell-specific expression of *Drosophila* IP_3_R^DN^ allowed the identification of two pairs of flight modulating dopaminergic neurons. Neuromodulatory signals received by these neurons stimulate IP_3_/Ca^2+^ signals to regulate critical aspects of pre-synaptic cellular physiology with significant impact on flight and feeding behaviour.

## RESULTS

### *Ex vivo* characterization of a dominant negative IP_3_R

To understand how Ca^2+^ release through IP_3_R affects cellular properties of neurons, we designed a mutant *itpr* cDNA to function as a dominant negative upon overexpression in wild type *Drosophila* neurons. The dominant negative construct (*Itpr*^*DN*^) was designed based on previous studies in mammalian IP_3_Rs (Figure 1A, B; Alzayady et al., 2016). Three conserved basic residues in the ligand binding domain (R272, K531 and Q533) of the *Drosophila* IP_3_R cDNA were mutated to Glutamine (Q) and the resultant mutant cDNA was used to generate GAL4/UAS inducible transgenic strains (*UASItpr*^DN^) as described in methods. Expression from the *Itpr*^*DN*^ construct was validated by western blots of adult fly head lysates. IP_3_R levels were significantly enhanced in fly heads with *Itpr*^DN^ as compared to genetic controls and were equivalent to overexpression of a wild-type IP_3_R transgene (*Itpr*^*+*^; Figure 1C). IP_3_-mediated calcium release from the IP_3_R in the presence of *Itpr*^*DN*^, was tested on previously characterised glutamatergic interneurons in the larval ventral ganglion known to respond to the muscarinic acetylcholine receptor (mAChR) ligand Carbachol. These neurons can be marked with a GAL4 strain (*vGlut*^*VGN6341*^; Jayakumar et al., 2016; 2018). Changes in cytosolic calcium were measured by visualising Ca^2+^ dependent fluorescence changes of a genetically encoded Ca^2+^ sensor GCaMP6m (Chen et al., 2013) in *ex vivo* preparations. *vGlut*^*VGN6341*^ marked neurons expressing IP_3_R^DN^ exhibit reduced as well as delayed Ca^2+^ responses to Carbachol stimulation as compared to controls (Figure 1D). Ca^2+^-released from the IP_3_R is also taken up by mitochondria (Bartok et al., 2019). Hence we measured mitochondrial Ca^2+^ uptake post Carbachol stimulation using mitoGCaMP, a mitochondrial targeted fluorescence sensor for Ca^2+^ (Lutas et al., 2012). Similar to cytosolic GCaMP, attenuated Ca^2+^ responses were observed upon carbachol stimulation in presence of IP_3_R^DN^ (Figure 1E). Interestingly, the mitochondrial response did not exhibit a delay in reaching peak values. We attribute the residual Ca^2+^ release observed in the cytosol and mitochondria to the persistence of some IP_3_R tetramers with all four wild type subunits, encoded by the native *itpr* gene. In summary, both cytosolic and mitochondrial Ca^2+^ measurements confirmed the efficacy of the *Drosophila* IP_3_R^DN^ for attenuating IP_3_-mediated Ca^2+^ release from ER stores upon GPCR stimulation in *Drosophila* neuron

**Figure 1:**
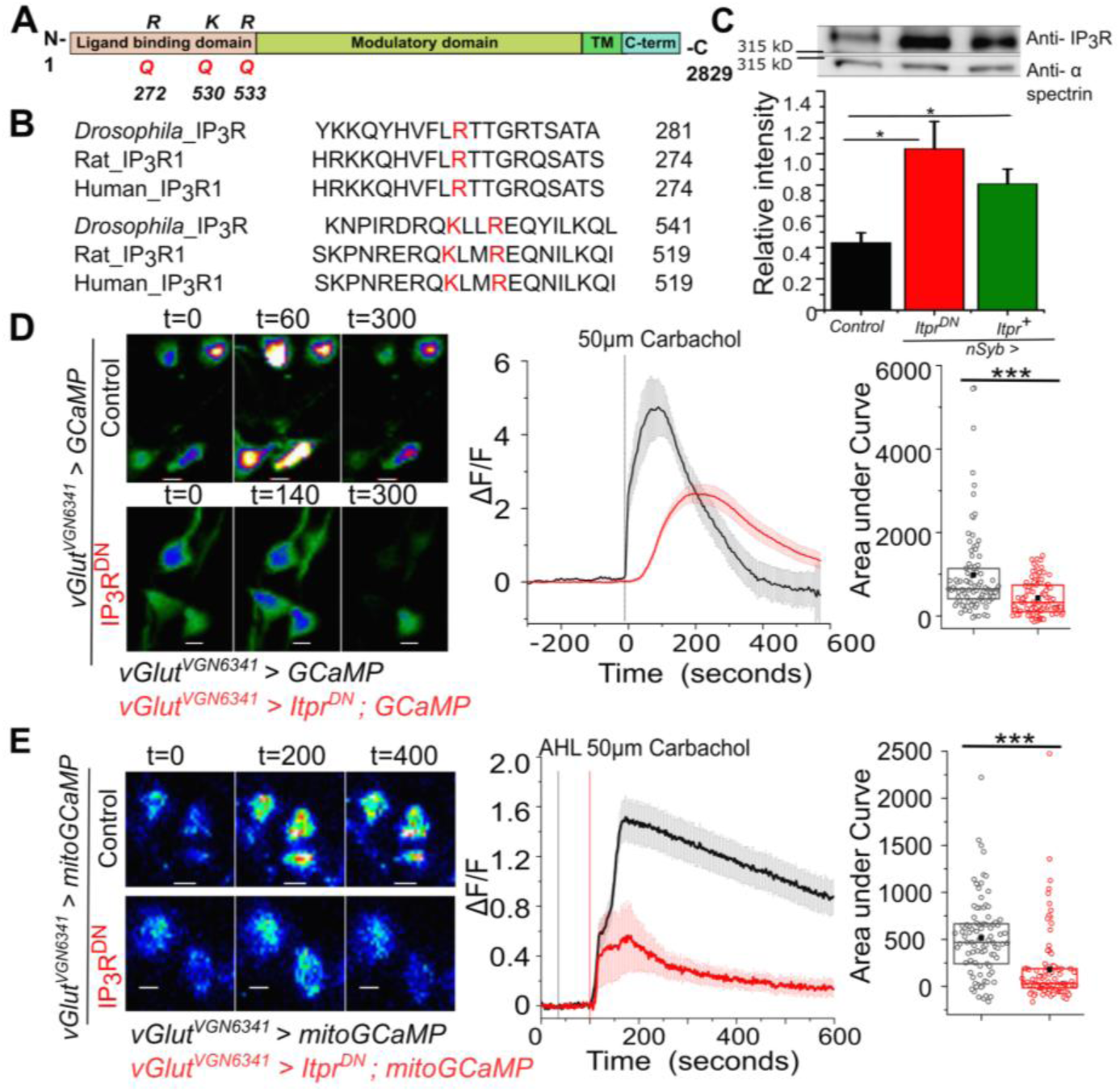
Generation and characterisation of a dominant negative IP_3_ R (IP _3_R^DN^) A. Domain organization of the *Drosophila* IP_3_R. The Arginine (R) and Lysine (K) residues mutated to Glutamine (Q) to generate a dominant-negative IP_3_R (IP_3_R^DN^) are shown. B. Alignment of the Drosophila IP_3_R with Rat IP_3_R1 and Human IP_3_R1 in the region of the mutated amino acids. All three residues (red) are conserved. C. Significantly higher immuno-reactivity against the IP_3_R is observed upon pan-neuronal (*nsybGAL4)* expression of IP_3_R^DN^ (*UASitpr*^*DN*^) and IP_3_R (*UASitpr*^*+*^) in adult heads. Statistical comparison was made with respect to *Canton S* as control, n=5, *p< 0.05 t-test. D. Representative images show changes in cytosolic Ca^2+^ in larval neurons of the indicated genotypes as judged by GCaMP6m fluorescence at the indicated time intervals after stimulation with Carbachol (Scale bars indicate 5 μm). Warmer colors denote increase in [Ca^2+^]_cyt_. Mean traces of normalized GCaMP6m fluorescence in response to Carbachol (4s/frame) in *vGlut*^*VGN6341*^*GAL4* marked neurons, with (red) and without (black) IP_3_R^DN^ (ΔF/F ± SEM) (middle panel). Area under the curve (right) was quantified from 0 - 420s from the traces in the middle. *vGlut*^*VGN6341*^*GAL4 > UAS GCaMP* N = 6 brains, 92 cells; *vGlut*^*VGN6341*^*GAL4 > UAS Itpr*^*DN*^; *UAS GCaMP* N = 6 brains, 95 cells. ***p< 0.005 (Two tailed Mann-Whitney U test). **E**. Representative images show changes in mitochondrial Ca^2+^ in larval neurons of the indicated genotypes as judged by mitoGCaMP6m fluorescence at the indicated time intervals after stimulation with Carbachol (Scale bars indicate 5 μm). Warmer colors denote increase in [Ca^2+^]_cyt_. Mean traces of normalized mitoGCaMP6m fluorescence in response to Carbachol (2s/frame) in *vGlut*^*VGN6341*^*GAL4* marked neurons, with (red) and without (black) IP_3_R^DN^ (ΔF/F ± SEM) (middle panel). Area under the curve (right) was quantified from 100 - 400s from the traces in the middle. *vGlut*^*VGN6341*^*GAL4 > UAS mitoGCaMP* N = 5 brains, 69 cells; *vGlut*^*VGN6341*^*GAL4 > UAS Itpr*^*DN*^; *UAS mitoGCaMP* N = 5 brains, 68 cells. ***p< 0.005 (Two tailed Mann-Whitney U test).

### The IP_3_R is required in a subset of central dopaminergic neurons for maintenance of *Drosophila* flight

Next we tested the functional efficacy of *Itpr*^*DN*^ for attenuating neuronal function in *Drosophila*. From previous reports we know that *itpr* mutants are flightless and their flight deficit can be rescued partially by overexpression of a wild-type cDNA construct (*Itpr*^*+*^; Venkatesh et al., 2001) in monoaminergic neurons (Banerjee, 2004). Subsequent studies identified mild flight deficits upon RNAi mediated knock down of the IP_3_R in dopaminergic neurons (DANs) (Pathak et al., 2015) when tethered flight was tested for 30 seconds. Knock-down of the IP_3_R in serotonergic neurons, that form another major subset of the tested monoaminergic neurons, did not give a flight deficit (Sadaf et al., 2012). Here we tested if pan-neuronal expression of the IP_3_R^DN^ affects longer flight bout durations in a modified tethered flight assay lasting for 15 mins (see methods and Manjila & Hasan, 2018). Flies with pan-neuronal *nsybGAL4* driven expression of *Itpr*^DN^ exhibit significantly reduced flight bouts (281.4±38.9s) as compared to the appropriate genetic controls *Itpr*^*DN*^*/+* (773.3±30.4s) and *nsyb/+* (670.6±41.3s) (Figure 2-figure supplement 1A). Stronger deficits in flight bout durations (185.2±33.7 seconds) were obtained by expression of *Itpr*^*DN*^ in Tyrosine Hydroxylase expressing cells, that include a majority of dopaminergic neurons (*THGAL4*; Friggi-Grelin et al., 2003), as well as in a dopaminergic neuron subset (260±30.5 s) marked by *THD’GAL4* (Liu et al., 2012) (Figure 2A, 2B). As additional controls the same GAL4 drivers were tested with overexpression of *Itpr*^*+*^. Interestingly, flight deficits were also observed upon overexpression of *Itpr*^*+*^ across all neurons (*nsyb>Itpr*^*+*^; 410±31.6s; Figure 2-figure supplement 1A), all TH expressing cells (*TH>Itpr*^*+*^; 482.2±41.4s) and a dopaminergic neuron subset, (*THD’>Itpr*^*+*^; 433.5±35.6s; Figure 2A), though these were milder than with expression of *Itpr*^DN^.

**Figure 2:**
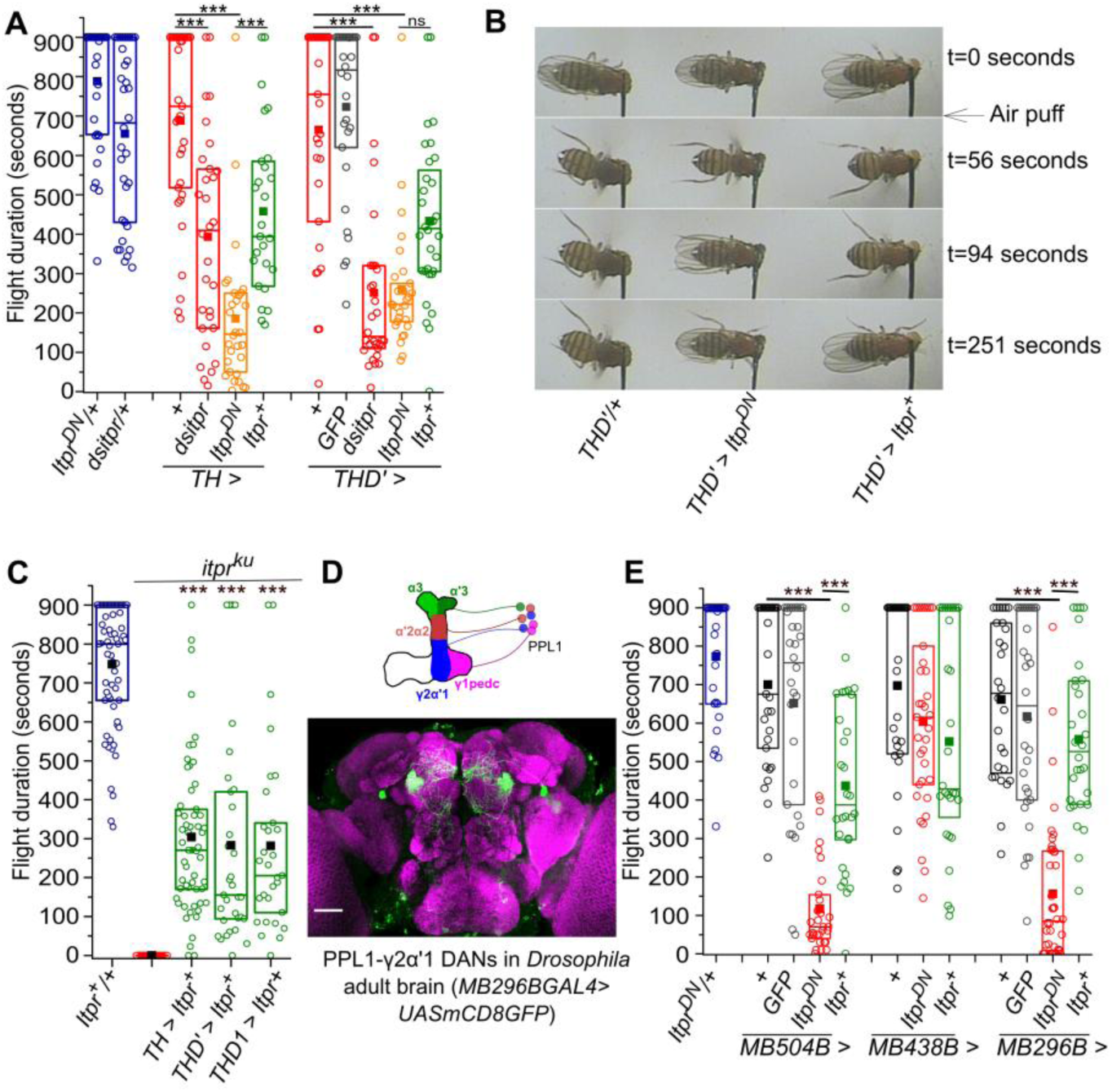
The IP_3_R is required in central dopaminergic neurons for maintaining long flight bouts. A. Flight deficits observed in flies expressing IP_3_ R RNAi (*dsitpr*), IP_3_ R^DN^ (*Itpr*^*DN*^) and IP_3_ R^WT^ (*Itpr*^*+*^) across all dopaminergic and TH expressing cells (*TH*) as well as a subset of dopaminergic neurons (*THD’*). Flight times of flies of the indicated genotypes are represented as box plots where the box represents 25-75% of the distribution, each circle is the flight time of an individual fly, the small filled square represents mean flight time and the horizontal line is the median. Flight was tested in flies with GFP in *THD’* marked neurons as an over-expression control. B. Snapshots from flight videos of air puff stimulated flight bouts in the indicated genotypes at the specified time points. C. Box plots (as in A above) represent flight bout durations of control and *itpr* mutant flies (*itpr*^*ku*^). Expression of a wild-type cDNA for the IP_3_R (*UAS Itpr*^*+*^) with indicated dopaminergic *GAL4s* rescued flight to a significant extent. D. A schematic of PPL1 DANs projecting to different lobes of one half of the mushroom body (top); An adult brain with GFP expression (green) driven by *MB296BGAL4*. GFP expression is restricted to PPL1-γ2α′1 neurons and the γ2α′1 MB lobe. The brain neuropil is immunostained with anti Brp (purple). Scale bar = 50 μm. E. Box plots (as in A) of flight bout durations in flies expressing IP_3_ R^DN^ (*Itpr*^*DN*^) and IP _3_R^WT^ (*Itpr*^*+*^) in the indicated *PPL1 DAN* splitGAL4 strains. In all box plots n > 25, ***p< 0.005 **p< 0.01, n.s., not significant at p < 0.05 by two-tailed Student’s t test (for C) or one-way ANOVA followed by post hoc Tukey’s test (for A and E). Comparisons for significance were with the control values except where marked by a horizontal line.

To confirm that shorter flight bouts in flies expressing *Itpr*^*DN*^ are a consequence of reduced IP_3_R function in dopaminergic neurons and not an overexpression artefact, we expressed a previously validated *Itpr* RNAi (Agrawal et al., 2010) and recorded flight durations in an identical flight assay. Significantly reduced flight bouts, comparable to the flight deficits obtained upon expression of *Itpr*^*DN*^, were observed by knockdown of the IP_3_R with *nsybGAL4, THGAL4* and *THD’GAL4* (Figure 2A, Figure 2-figure supplement 1A, Figure 2-figure supplement 2). These data confirm that IP_3_R function is necessary in the *THD’* marked subset of dopaminergic neurons for maintenance of flight bouts. Further, it suggests that either decrease (*Itpr*^*DN*^ and *Itpr RNAi*) or increase (*Itpr*^+^) of IP_3_-mediated Ca^2+^ release in *THD’* neurons affects flight bout durations.

As an independent test of IP_3_R requirement, the wild-type IP_3_R was overexpressed in dopaminergic neurons of an adult viable IP_3_R mutant, *itpr*^*ka1091/ug3*^ or *itpr*^*ku*^ (Joshi et al., 2004). Overexpression of the IP_3_R in monoaminergic neurons of *itpr* mutants can rescue free flight measured for short durations of 5-10 secs (Banerjee, 2004). Short (30s, Figure 2-figure supplement 1B) and long (900s; Figure 2C) flight bouts were measured in a heteroallelic viable *itpr* mutant combination *itpr*^*ku*^, with *Itpr*^+^ overexpression in all neurons or in dopaminergic neurons and dopaminergic neuronal subsets. Pan-neuronal overexpression of the IP_3_R (*nsyb>Itpr*^*+*^) rescued short flight bouts partially in 10 out of 30 flies tested (Figure 1-figure supplement 1B). Interestingly, complete rescue of short flight was observed in flies rescued by IP_3_R overexpression in dopaminergic neurons and their TH-D subsets but not the TH-C’ subset (Figure 2 - figure supplement 1B). Better rescue by IP_3_R overexpression in dopaminergic neurons suggests weak expression of *nSybGAL4* in dopaminergic neurons, though this idea needs further verification. When tested for longer flight bouts, a partial rescue from dopaminergic neurons and TH-D subsets was observed (304.2±25.1s; *TH>Itpr+*) as compared to control flies (747.8±19.4s; Figure 2C), suggesting that the IP_3_R regulates flight bout durations from both dopaminergic and certain non-dopaminergic neurons (Agrawal et al., 2010). The *TH-D* and *TH-C GAL4*s express in anatomically distinct central dopaminergic neurons of which the *THD1GAL4* and *THD’GAL4* uniquely mark the PPL1 and PPM3 DAN clusters. Taken together, shorter flight bouts in *THD>Itpr*^*DN*^ flies and significant rescue of flight by *THD>Itpr*^*+*^ overexpression in flightless *itpr*^*ku*^ identifies an essential requirement for IP_3_R function in the PPL1 and/or PPM3 DANs for flight bouts lasting upto ∼300s.

The two PPL1 clusters on each side of the brain consist of 12 pairs of neurons (Mao & Davis, 2009) and have been implicated in the maintenance of long flight bouts previously (Pathak et al., 2015). To further restrict flight modulating neuron/s in the PPL1 group we identified split*GAL4* strains that mark fewer PPL1 neuron/s and project to individual lobes of the Mushroom Body (MB; Figure 2-figure supplement 1C and Figure 2D) (Aso et al., 2014; Aso & Rubin, 2016). Amongst the identified split*GAL4* strains, flight deficits were observed by expression of IP_3_R^DN^ in PPL1 DANs projecting to the MB lobes α’2α2, α3, γ1peduncle and γ2α′1 (*MB504BGAL4*), but not with PPL1 DANs projecting to the MB lobes α’2α2, α3 and Y1peduncle (*MB438BGAL4*; Figure 2E), suggesting PPL1-γ2α′1DANs as the primary focus of IP_3_R function. Indeed, expression of *Itpr*^*DN*^ in PPL1-γ2α′1 DANs, marked by *MB296BGAL4*, resulted in significantly shorter flight bouts, when compared to *Itpr*^*+*^ expression in the same cells (Figure 2E). Expression of *Itpr*^*DN*^ in other PPL1 neurons failed to exhibit significant flight deficits (Figure 2 - figure supplement 1D). Thus *PPL1-γ2α′1* DANs require IP_3_R function to sustain longer flight bouts. These data do not exclude a role for the IP_3_R in PPM3 DANs that are also marked by *THD’GAL4* in the context of flight.

**Figure 2-figure supplement 1:**
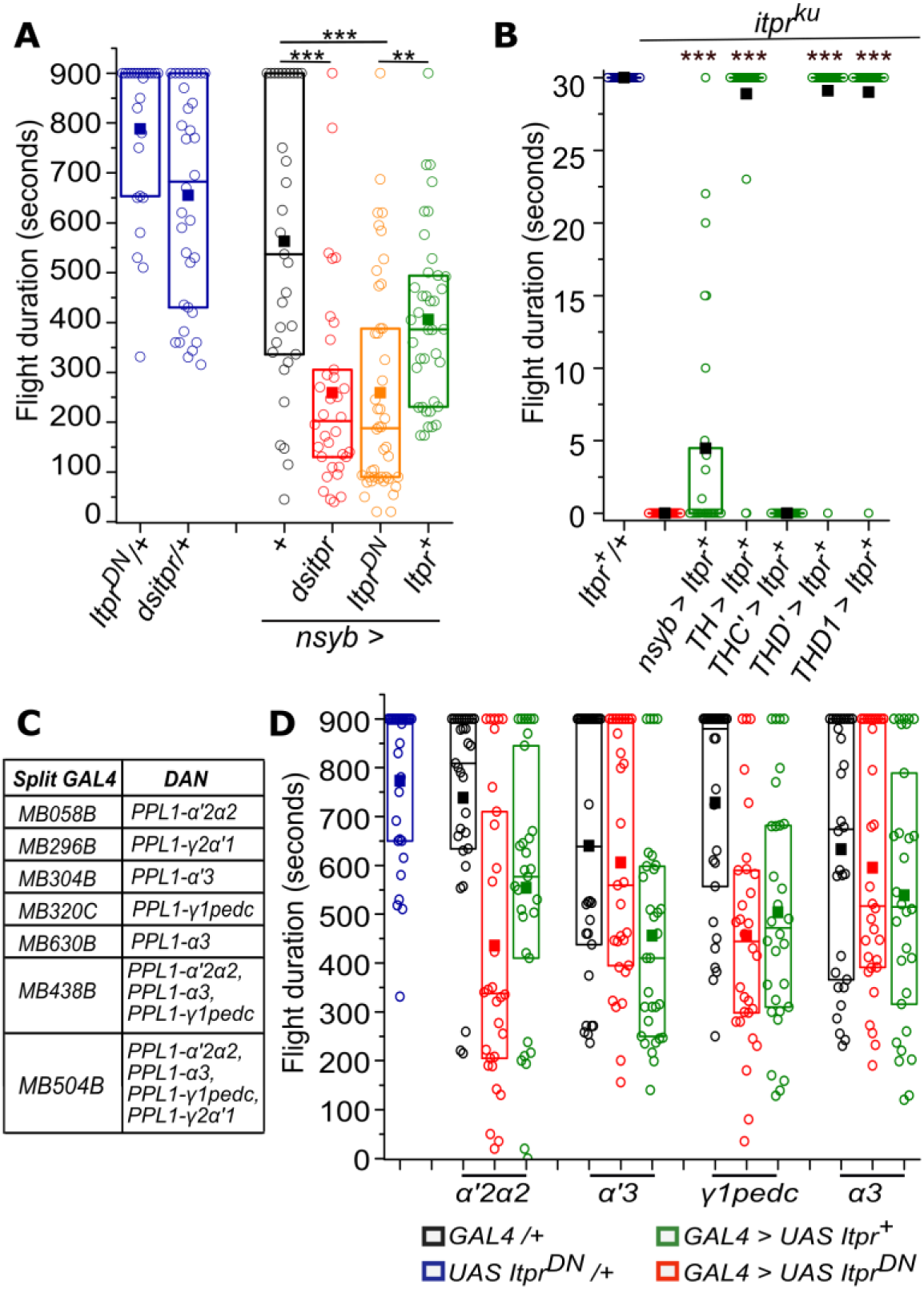
Perturbation of IP_3_R signaling in neurons affects the duration of flight bouts. **A)** Box plots represent flight deficits observed upon perturbing IP_3_R signaling after expressing IP_3_R RNAi (*dsitpr*), IP _3_ R^DN^ (*Itpr*^*DN*^) and IP_3_ R^WT^ (*Itpr*^*+*^) across all classes of neurons with *nsybGAL4*. **B)** Box plots represents flight bout durations of *itpr* mutant flies measured for 30 seconds, upon expression of *UAS Itpr*^*+*^ with the indicated *GAL4s*. **C)** SplitGAL4 strains used to label subsets of PPL1 DANs. **D)** Box plots represent flight bout durations of flies upon perturbing IP_3_ R signaling after expressing IP_3_ R^DN^ (*Itpr*^*DN*^) and IP_3_ R^WT^ (*Itpr*^*+*^) in the indicated *PPL1 DAN* expressing strains. Box plot symbols are as described in methods and Figure 2A; n ≥ 30, ***p< 0.005 **p< 0.05, n.s., not significant at p < 0.05 by two-tailed Student’s t test (for B) or one-way ANOVA followed by post hoc Tukey’s test (for A and D. All comparisons for significance were with the control values except where marked by a horizontal line. Comparisons for significance were with *itpr* mutants in B.

**Figure 2-figure supplement 2:** Video showing flight defect in *THD’> Itpr*^*DN*^ (Right) as compared to control (*THD’/+*, Left)

### The IP_3_R is required in a late developmental window for adult flight

Shorter flight bouts in adults might arise either due to loss or change in properties of identified neurons during development. Alternately, the IP_3_R might acutely affect the function of these neurons during flight in adults. To distinguish between these possibilities we employed the TARGET system (McGuire et al., 2003) for temporal control of IP_3_R^DN^ expression in *PPL1* and *PPM3* DANs. TARGET uses a ubiquitously expressed temperature sensitive repressor of GAL4 (Tub-GAL80^ts^) that allows GAL4 expression at 29°, but not at lower temperatures. Hence expression of a GAL4 driven *UAS* transgene can be controlled by changing incubation temperatures from 18° to 29°. *THD’GAL4* driven *Itpr*^*DN*^ expression was restricted to the larval stages (data not shown), 0-48h after puparium formation (APF), 48-96h APF and for 0-3 days in adults. Flight deficits were observed upon expression of IP_3_R^DN^ during pupal development and not when expressed in adults. In pupae the deficit was most prominent by expression of the *Itpr*^*DN*^ during the shorter window of 48-96hr APF (Figure 3A).

**Figure 3:**
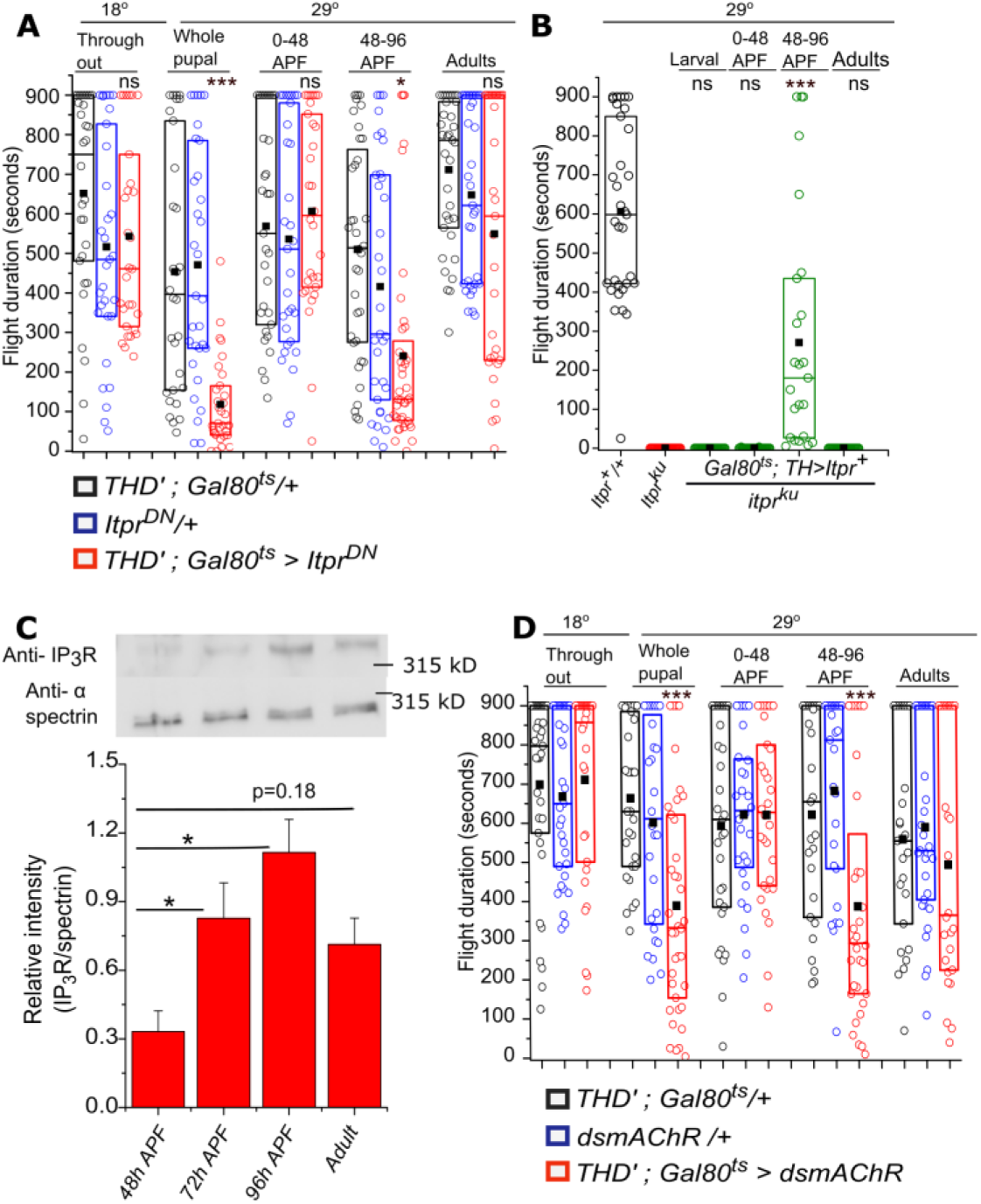
Adult flight phenotypes arise from late pupal expression of the IP_3_R and mAChR. A) Box plots of flight bout durations in adults after temporal expression of IP_3_R^DN^ in *THD’* neurons (*THD’; TubGAL80*^*ts*^ > *Itpr*^*DN*^) by transferring them to 29°C at the indicated stages of development. For adult expression, flies were transferred to 29°C immediately after eclosion and tested for flight after 3 days. B) Box plots of flight bout durations after expressing the IP_3_R (*Itpr*^*+*^) in dopaminergic cells of *itpr* mutants at the indicated stages of development. For stage specific expression a *TH; TubGAL80*^*ts*^ strain was used and the progeny transferred to 29°C at appropriate time developmental points. C) Levels of the IP_3_R increase in late pupae between 72-96 hrs APF and plateau between 96hrs APF and adults (3 days). A representative western blot from lysates of dissected central nervous systems of *Canton S* probed with anti-IP_3_R and anti-spectrin is shown (top). Three independent lysates and blots were quantified (below). *p< 0.05, t-test. D) Box plots with flight bout durations of adult flies after knockdown of the Muscarinic acetylcholine receptor (mAChR) in *THD’* neurons (*THD’;TubGAL80*^*ts*^ > *dsmAChR*) at the indicated stages of pupal development and in adults. Box plot symbols are as described in methods, n ≥ 30, ***p< 0.005 **p< 0.05, n.s., not significant at p < 0.05 by one-way ANOVA followed by post hoc Tukey’s test (for A and D) and n ≥ 23, ***p< 0.005, n.s., not significant at p < 0.05 by two-tailed Student’s t test (for B). All comparisons for significance were with the control values for A and C, with *itpr* mutants for B.

Requirement for the IP_3_R during the 48-96hr APF window was independently confirmed by measuring flight after temporal expression of *Itpr*^*+*^ in dopaminergic neurons of *itpr*^*ku*^ (Figure 3B). Flight in *itpr*^*ku*^ mutant animals was rescued to a significant extent by *Itpr*^*+*^ expression from 48-96h APF (277±60.3s) but not when expressed before and after. Interestingly, post-48 hr APF is also the time period during which levels of the IP_3_R are upregulated in the pupal brain (Figure 3C). Thus expression from the *Itpr*^*DN*^ encoding transgene during the 48-96hr interval presumably results in the formation of a majority of inactive IP_3_R tetramers due to the presence of at least one IP_3_R^DN^ monomer. The cellular and physiological consequences of inactive IP_3_Rs in pupal, and subsequently adult brains, was investigated next.

It is known that specification of central dopaminergic neurons is complete in late larval brains (Hartenstein et al., 2017) The number of PPL1 DANs was no different between controls and in flies expressing *Itpr*^*DN*^ in either *TH-D* or *MB296B* marked neurons (Figure 3 - figure supplement 1A). This finding agrees with our observation that larval expression of *Itpr*^*DN*^ had no effect on adult flight. Moreover projections from *MB296B* to the *γ2α′1* MB lobes also appeared unchanged upon expression of *Itpr*^*DN*^ (Figure 3 - figure supplement 1B).

To understand the nature of signaling through the IP_3_R, required during pupal development we tested flight after knockdown of an IP_3_/Ca^2+^ linked GPCR, the muscarinic acetylcholine receptor (mAChR). From a previous study it is known that neuronal expression of the mAChR during pupal development is required for adult flight (Agrawal et al., 2013). Adult flight bouts were significantly shorter upon stage specific knock-down (48-96h APF) of the mAChR in TH-D’ neurons with an RNAi under temporal control of the TARGET system (Figure 3D). Further knockdown of mAChR in PPL1-*γ2α′1* DANs also manifested mild flight defect (Figure 3 - figure supplement 1C). These data suggest that the mAChR and IP_3_-mediated Ca^2+^ release are required during pupal maturation of a central brain circuit that functions for the maintenance of long flight bouts. Alternately, the pupal requirement may arise from the fact that both the IP_3_R (Figure 3A, B) and the mAChR (Figure 3D) are synthesized in late pupal neurons, carried over to adult neurons where they have a slow turnover, and function during acute flight. Taken together our data support the idea that acetylcholine, a neurotransmitter, activates the mAChR on PPL1 DANs of late pupae and/or adults to stimulate Ca^2+^ release through the IP_3_R. Cellular changes arising from loss of IP_3_ mediated Ca^2+^ release were investigated next.

**Figure 3 - figure supplement 1:**
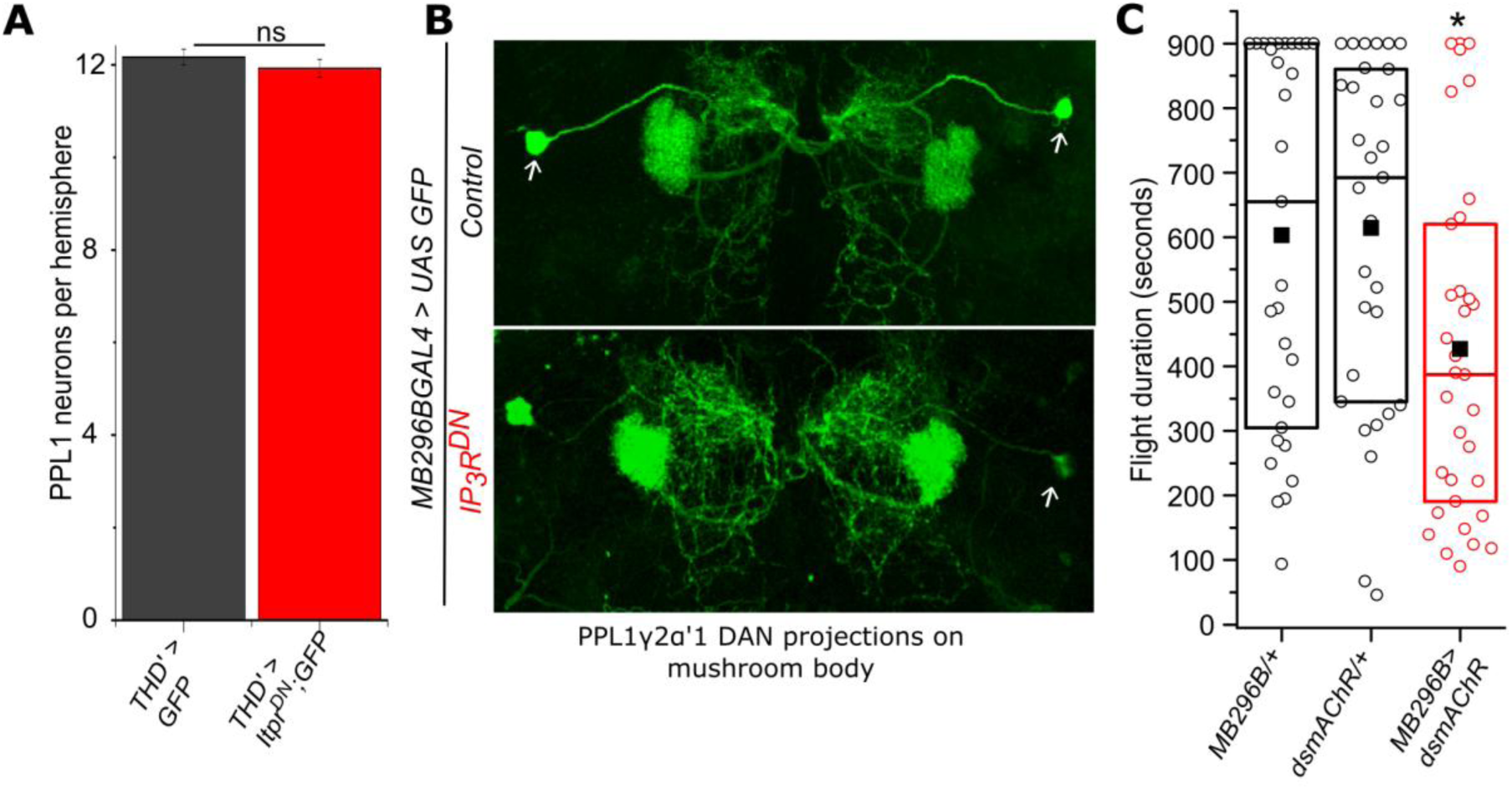
Expression of IP_3_R^DN^ and mAChR RNAi in *THD’* and *MB296B* marked neurons. A) Expression of IP_3_R^DN^ does not change the number of PPL1 neurons. Cells were quantified from both hemi-segments of N=6 brains. B) PPL1-γ2α′1 neurons marked by *MB296BGAL4* are shown (white arrows) with their pattern of innervation to the MB neuropil as visualized using membrane-localized GFP. Expression of IP_3_R^DN^ (lower panel) did not affect the pattern of MB innervation. C) Box plots of flight bout durations in control flies and flies with knockdown of mAChR in *MB296B* cells. n ≥ 30, *p< 0.05 at p < 0.05 by one-way ANOVA followed by post hoc Tukey’s test

### Synaptic vesicle release and IP_3_R are both required for the function of *PPL1-γ2α′1* DANs

The 48-96 hr time window of pupal development is when adult neural circuits begin to mature with the formation of synapses, some of which are eliminated whilst others are strengthened (Akin et al., 2019; Consoulas et al., 2002; K. X. Zhang et al., 2016). Synapse strengthening occurs when pre-synaptic neurotransmitter release leads to post-synaptic excitation/inhibition (Andreae & Burrone, 2018; Baines, 2003; Baines et al., 2001; Pang et al., 2010). We hypothesized that IP_3_-mediated Ca^2+^ release might modulate synaptic activity and hence lead to strengthening of synapses between *THD’* DANs and their post-synaptic partners during pupal development. To test this idea, the requirement for synaptic vesicle recycling in *THD’* and *PPL1-γ2α′1* DANs was investigated through pupal development. A transgene encoding a temperature sensitive mutant of Dynamin, *Shibire*^*ts*^ (Vicario et al., 2001), that prevents synaptic vesicle recycling at 29°C and hence blocks neurotransmitter release, was expressed during pupal development, by transfer to 29°C at the appropriate time interval. Loss of synaptic vesicle recycling in pupae resulted in significantly shorter flight bout durations of 419.9±50.2s (*THD’*) and 411.5±57s (*MB296B*) as compared to controls (Figure 4A and Figure 4 - figure supplement 1A). The requirement for synaptic vesicle release was further restricted to 48-96 hrs APF consistent with the requirement of IP_3_R at the same time interval (Figure 3A,B). These data suggest that *THD’* marked dopaminergic neurons require both synaptic vesicle recycling and IP_3_-mediated Ca^2+^ release during late pupal development for their adult function.

**Figure 4:**
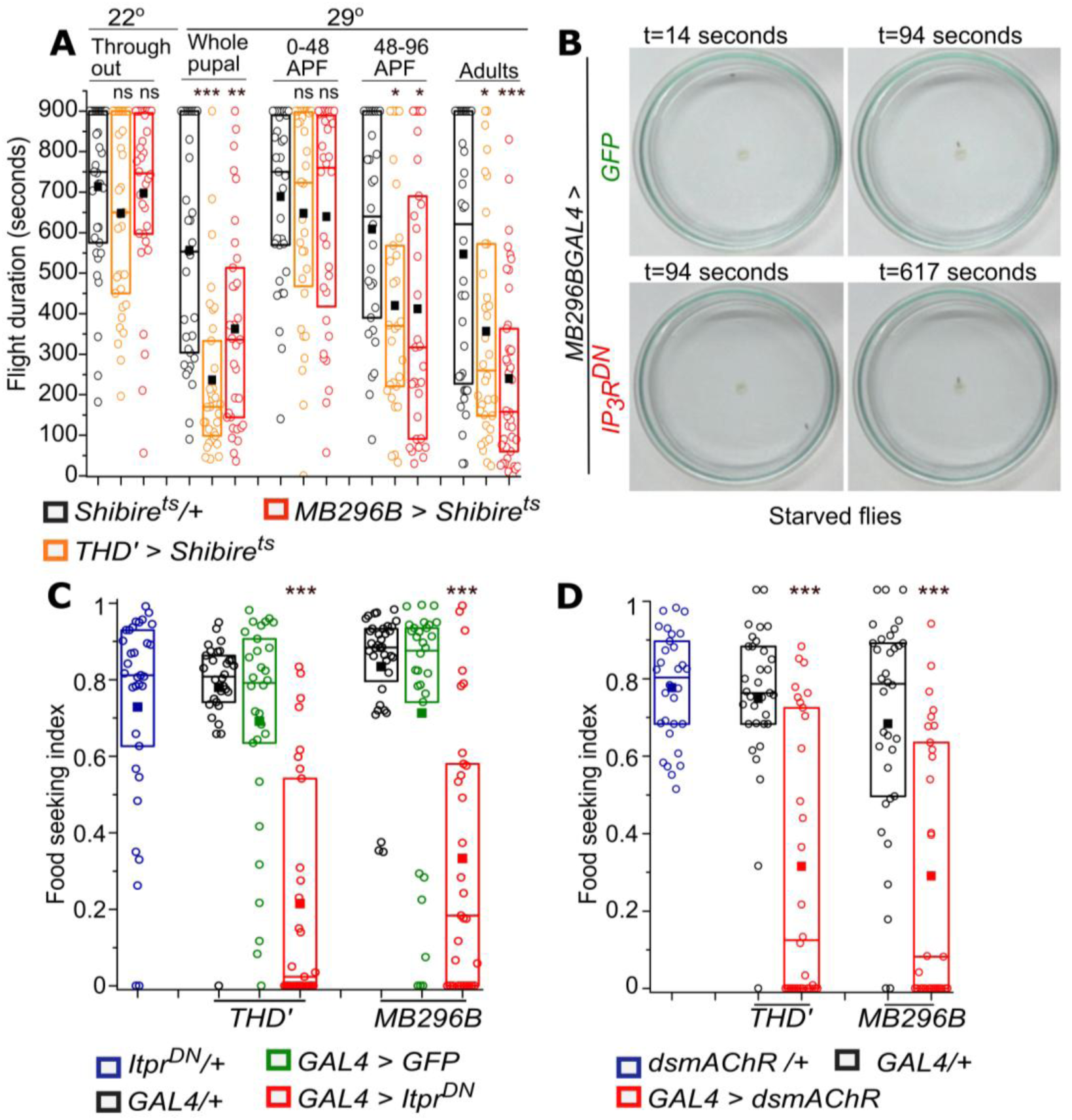
Synaptic vesicle recycling and IP_3_R function are required in PPL1 dopaminergic neurons for adult flight and feeding. A) Quantification of flight deficits observed upon blocking synaptic vesicle recycling by expression of a temperature sensitive Dynamin mutant, *Shibire*^*ts*^ in *THD’* and *MB296B* cells. Inactivation of Dynamin was achieved in strains expressing *Shibire*^*ts*^, by transfer to 29°C, at the indicated stages of development. For adults, flies were grown at 22°C and transferred to 29°C 10 minutes before the flight assay also performed at 29°C. n ≥ 30, ***p< 0.005, **p< 0.01, *p< 0.05 at p < 0.05 by one-way ANOVA followed by post hoc Tukey’s test. B) Food seeking by hungry flies is significantly diminished upon expression of IP_3_R^DN^ in PPL1-γ2α′1 DANs (*MB296B*). Snapshots at the indicated time points from videos of starved male flies of the indicated genotype seeking a drop of yeast placed in the centre of a petriplate C) Quantification of food seeking behavior in starved males of the indicated genotypes. Expression of IP_3_R^DN^ in *THD’* and *MB296B* cells reduced food seeking behavior to a significant extent (red). Expression of *GFP* did not affect the behavior (green). n ≥ 30, ***p< 0.005 at p < 0.05 by two tailed Mann-whitney U test. D) Quantification of food seeking behavior in starved males of the indicated genotypes. Knockdown of mAChR in *THD’* and *MB296B* cells reduced food seeking behavior to a significant extent (red, n ≥ 30, ***p< 0.005 at p < 0.05 by two tailed Mann-Whitney U test).

The acute requirement for synaptic vesicle recycling in adult PPL1 (*THD’*) and *PPL1-γ2α′1* subset (*MB296B*) of DANs for maintenance of flight bouts was tested next. A previous report shows that synaptic vesicle recycling is required in PPL1 DANs marked by *THD1GAL4* during active flight (Ravi et al., 2018). We tested if synaptic vesicle release is also required in *THD’* and *MB296B* marked PPL1 DANs in adults. Importantly, flight bout durations were reduced significantly after acute inactivation (5 min) of synaptic vesicle recycling in PPL1 DANs (*THD’*) and the pair of *PPL1-γ2α′1* subset (*MB296B*) DANs in adults, supporting a requirement for neurotransmitter release from these neurons during flight (Figure 4A).

Impaired synaptic vesicle release from adult *PPL1-γ2α′1 DANs*, by expression of Shibire^ts^, also reduces the ability of starved flies to identify a food source rapidly (Tsao et al., 2018). To undertand if the IP_3_R affects the function of *PPL1-γ2α′1* DANs in more than one behavioural context, we tested food seeking behaviour of starved flies expressing *Itpr*^*DN*^ in PPL1 (*THD’*) and *PPL1-γ2α′1* subset (*MB296B*) *DANs*. The food seeking index of hungry flies with IP_3_R^DN^ was reduced significantly as compared to controls (Figure 4B, 4C, Figure 4-figure supplement 2 and Figure 4-figure supplement 3). Food-seeking behaviour of fed flies of all genotypes tested appeared similar (Figure 4-figure supplement 1B).

Next we tested if knockdown of the mAChR, the identified flight regulating GPCR (Figure 3 and Figure 3 - figure supplement 1C) that couples to IP_3_/Ca^2+^ signaling, also affected food seeking in starved flies. Knockdown of mAChR in either PPL1 (*THD’*) or *PPL1-γ2α′1* decreased the ability of starved males to find a yeast drop (Figure 4D), while fed flies were similar to genetic controls (Figure 4 - figure supplement 1C). Thus cholinergic stimulation of IP_3_/Ca^2+^ in *PPL1-γ2α′1* DANs reduces the motivation for longer flight bouts as well as the motivation to search for food.

Taken together these data demonstrate a requirement for both synaptic vesicle recycling and IP_3_-mediated Ca^2+^ release in the modulation of behaviour by *PPL1-γ2α′1* DANs. Furthermore, they suggest that the IP_3_R might regulate synaptic function in *PPL1-γ2α′1* DANs.

**Figure 4 - figure supplement 1:**
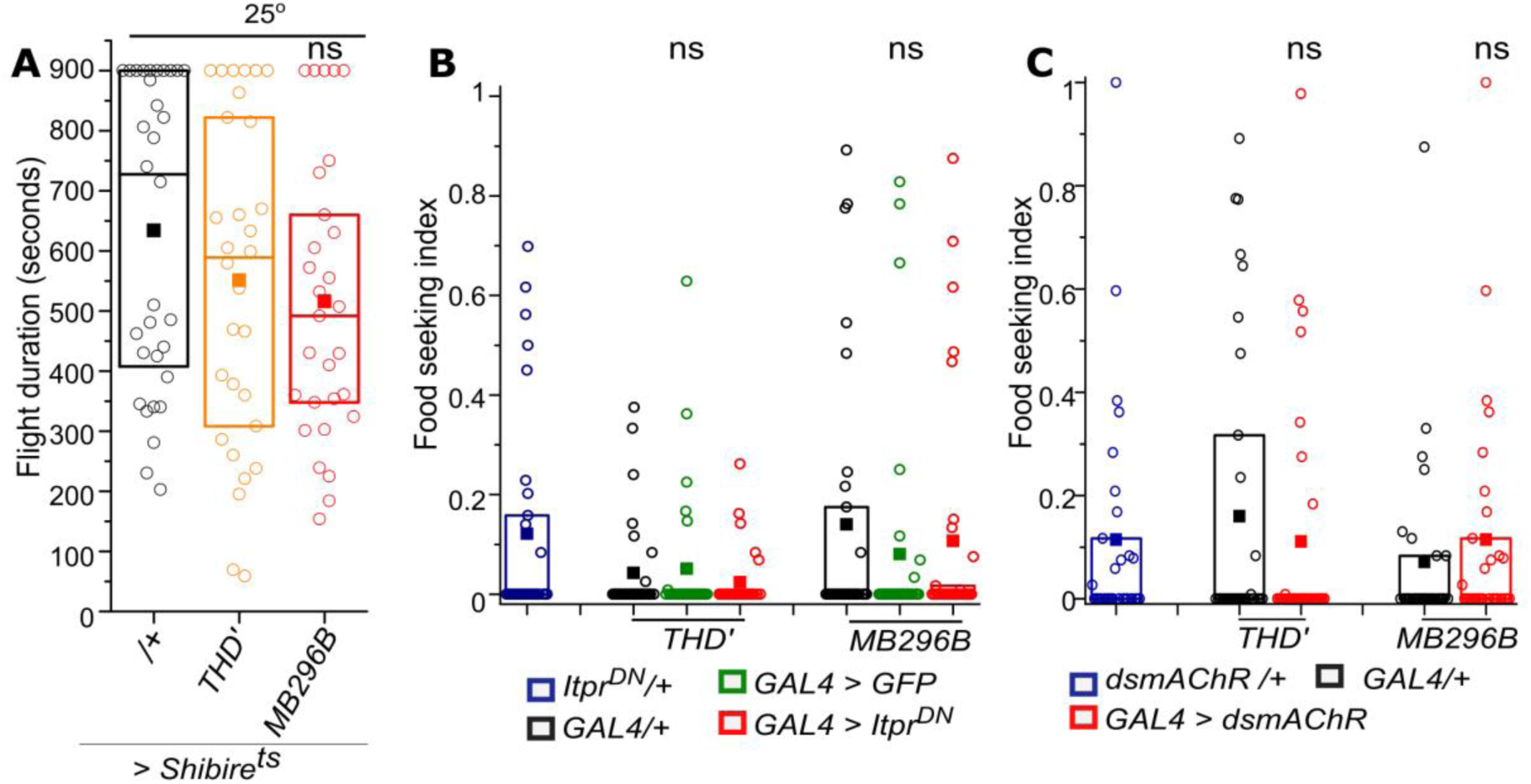
Flight is unaffected and food seeking behaviour is normal in control genotypes and conditions. A) Quantification of flight bouts upon expressing temperature sensitive dynamin mutant *Shibire*^*ts*^ in *THD’* and *MB296B* cells when flies were grown at the permissive temperature of 25°C. n ≥ 30, n.s., not significant at p < 0.05 by one-way ANOVA followed by post hoc Tukey’s test B) Food seeking behavior is absent in fed flies including the indicated control strains and upon expression of IP_3_ R^DN^ in *THD’* and *MB296B* cells (red). n ≥ 30, n.s., not significant at p < 0.05 by two tailed Mann-whitney U test. C) Food seeking behavior is absent in fed flies including the indicated control strains and upon knockdown of mAChR in *THD’* and *MB296B* cells (red). n ≥ 30, n.s., not significant at p < 0.05 by two tailed Mann-Whitney U test. **Figure 4 - figure supplement 2:** Video showing food seeking behavior in control fly (*MB296BGAL4/+*) **Figure 4 - figure supplement 3:** Video showing defect in food seeking behavior upon expression of IP_3_R^DN^ in *MB296B* cells.

### The IP_3_R affects neurotransmitter release from adult dopaminergic neurons

Decreased synaptic activity upon IP_3_R^DN^ expression might be a consequence of a reduction in synapse number. Therefore, a pre-synaptic marker, synaptotagmin tagged to GFP (*syt*.*eGFP;* Y. Q. Zhang et al., 2002), that localises to synaptic vesicles was expressed in *THD’* marked neurons in the absence and presence of IP_3_R^DN^. Fluorescence of syt.eGFP in the MB lobes was no different between control brains and in presence of the IP_3_R^DN^ (Figure 5A), suggesting that synapse numbers were unchanged by expression of IP_3_R^DN^.

**Figure 5:**
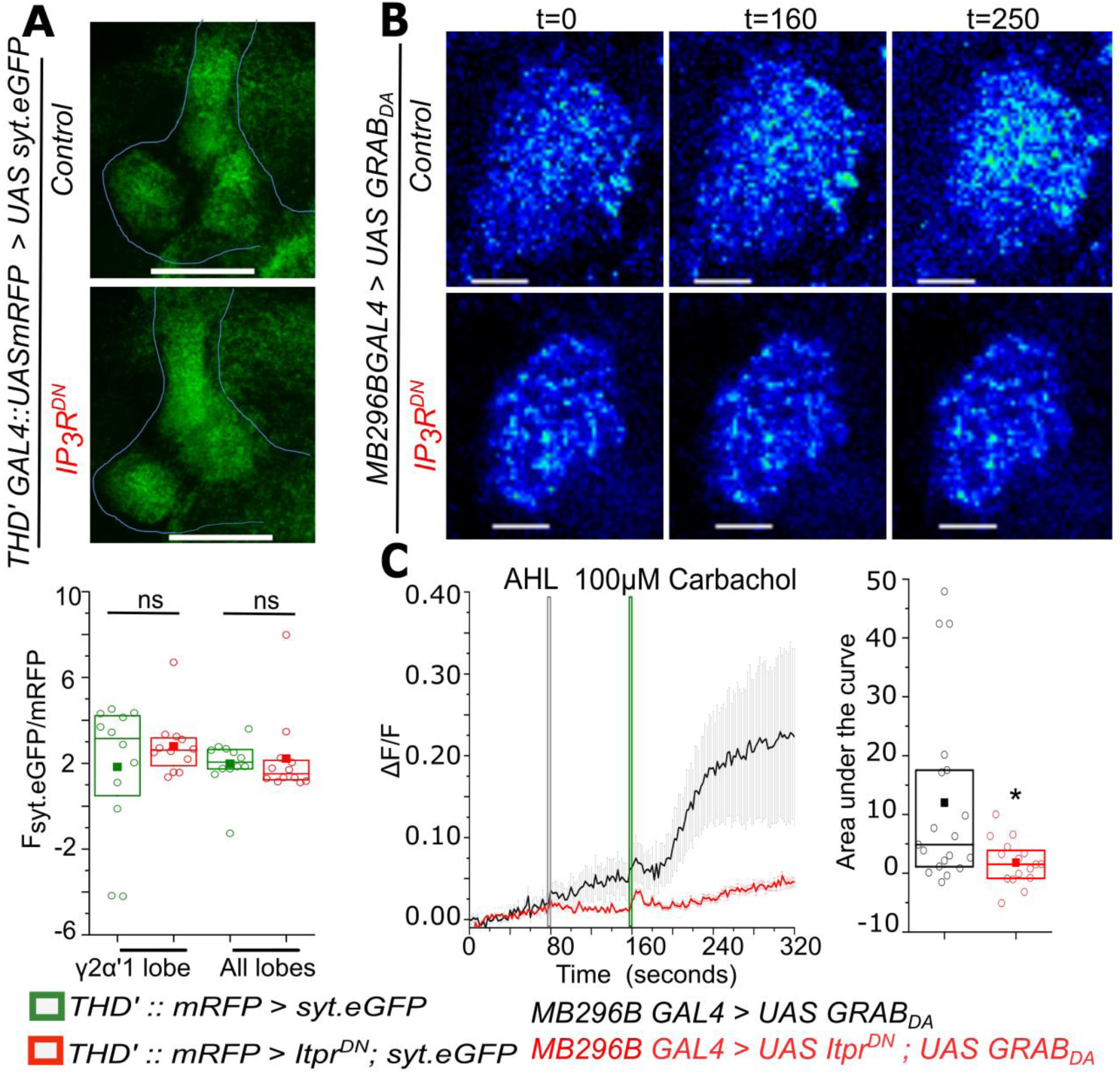
Carbachol-stimulated Dopamine release at the axonal terminals of *PPL1-γ2α′1* neurons is attenuated by expression of IP_3_ R^DN^. A) Representative images of a presynaptic marker synaptotagmin GFP (syt.eGFP) in lobes of the Mushroom body (top). Quantification of syt.eGFP fluorescence normalized to mRFP is shown below in the indicated MB regions marked by *THD’GAL4*. N = 6 brains, n.s. not significant, by Mann Whitney U test. Scale bars indicate 50 μm B) Carbachol stimulated dopamine release at PPL1-γ2α′1 termini visualized by changes in GRAB_DA_ fluorescence in a representative mushroom body lobe from brains of the indicated genotypes. Images were acquired at 2s per frame. Brighter fluorescence denotes increase in Dopamine. Scale bars = 10 μm **C)** Average traces (± SEM) of normalized GRAB_DA_ fluorescence responses (ΔF/F) upon addition of Carbachol (Green line) (panel on the left). Images were acquired at 2s per frame; Quantification of area under the curve was from the point of stimulation at 160s and up to 250 s. Boxplots and symbols are as described in Figure 2A. *MB296B GAL4 > UAS GRAB*_*DA*_ N = 10 brains,19 cells; *MB296B GAL4 > UAS Itpr*^*DN*^; *UAS GRAB*_*DA*_ N = 8 brains,16 cells. *p< 0.05, Mann-Whitney U test.

Neurotransmitter release from *PPL1-γ2α′1* (*MB296B*) DANs in response to stimulation of the IP_3_R was investigated next. Presence of the mAChR was confirmed on *PPL1-γ2α′1* DANs in adults by measuring Ca^2+^ release upon stimulation with the mAChR agonist Carbachol. Changes in GCaMP6m fluorescence in *MB296B* marked neurons post-Carbachol stimulation are shown in Figure 5 - figure supplement 1A. The ability of Carbachol to stimulate dopamine release from *MB296B* marked neurons in the *γ2α′1* MB-lobe was measured next. For this purpose a recently designed fluorescent sensor for dopamine, GRAB_DA_ (G protein-coupled receptor-activation based DA sensor; Sun et al., 2018) was expressed in *MB296B* neurons followed by stimulation with Carbachol. GRAB_DA_ consists of a dopamine receptor linked to cpEGFP such that its fluorescence increases upon binding of dopamine. Thus, a qualitative measure of dopamine release at the synaptic cleft is the change in GRAB_DA_ fluorescence when recorded in the *γ2α′1* MB lobe, the site of *PPL1-γ2α′1* synapses (Figure 5B). Control brains, with detectable changes in GRAB_DA_ fluorescence (ΔF/F) above an arbitrary value of 0.05 within 140 seconds of Carbachol stimulation, were classified as responders. Responding brains decreased from 77% (10/13 brains) in controls to 47% (8/17 brains) upon expression of IP_3_R^DN^. Moreover, GRAB_DA_ fluorescence arising from dopamine release was significantly muted amongst responders expressing the IP_3_R^DN^ as compared to responders from control brains (Figure 5B and Figure 5-figure Supplement 1B). Thus expression of IP_3_R^DN^ in *MB296B* DANs reduced the synaptic release of dopamine at the *γ2α′1* lobe. Interestingly, increase in GRAB_DA_ fluorescence at the synaptic terminals in the MB was faster (Figure 5C) than the carbachol-stimulated change in GCaMP fluorescence measured in the cell body (Figure 5-figure Supplement 1A). Perhaps there exist a greater number of mAChRs at the synapse than on the soma of *MB296B* DANs,as mAChRs are shown to be present on Kenyon cells and mushroom body (Bielopolski et al., 2019; Kondo et al., 2020). Though, requirement for the IP_3_R and the mAChR was restricted to 48-96 hrs APF, these data suggest that the neurotransmitter acetylcholine stimulates dopamine release in adult DANs through mAChR-IP_3_R signaling. The lack of flight deficits by adult-specific expression of IP_3_R^DN^ (Figure 3A) and knockdown of mAChR (Figure 3C) is probably due to perdurance of the respective proteins from late pupae through to adults.

**Figure 5 - figure supplement 1:**
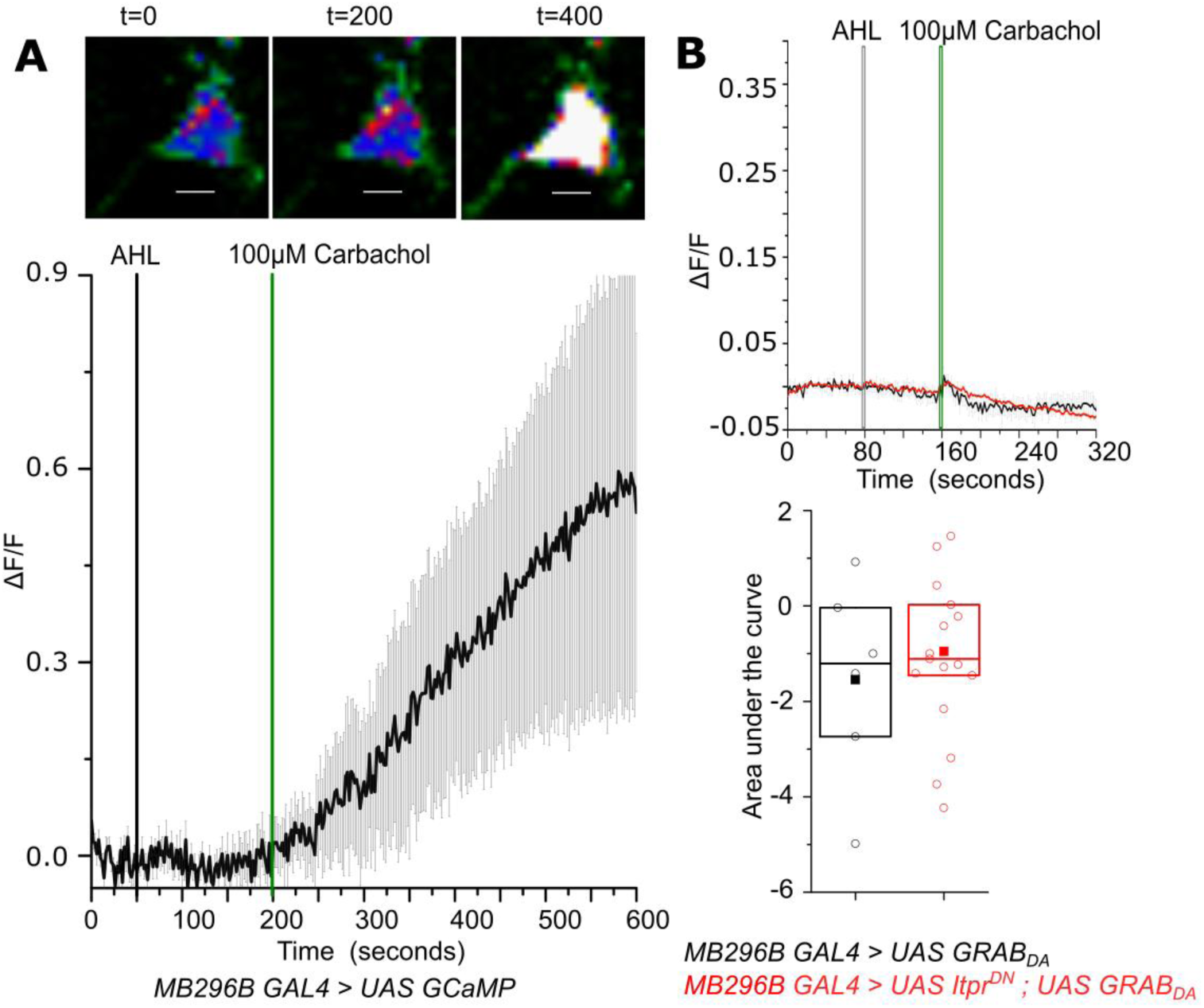
Carbachol evoked Ca^2+^ responses in the soma of PPL1-γ2α′1 DANs marked by *MB296BGAL4*. A) Carbachol stimulated change in GCaMP6m fluorescence (ΔF/F) in PPL1-γ2α′1 cells. Representative images of GCaMP6m fluorescence at the indicated time points (top panel). Warmer colors denote increase in [Ca^2+^]. Average response (± SEM) of normalized changes in GCaMP6m fluorescence (ΔF/F) in *MB296BGAL4* marked PPL1 cells upon addition of Carbachol from N=7 brains, 12 cells. Images were acquired at 2s per frame B) Average traces (± SEM) of normalized changes in GRAB_DA_ fluorescence (ΔF/F) from axonal termini of *MB296B*GAL4 marked DANs that responded below the arbitrary threshold (ΔF/F=0.05) upon addition of Carbachol (Green line). Images were acquired at 2s per frame (Top panel). Quantification of area under the curve (AUC) from the point of stimulation 160s up to 250s. Boxplots and symbols are as described in Figure 2A. *MB296B GAL4 > UAS GRAB*_*DA*_ N = 3 brains, 6 cells; *MB296B GAL4 > UAS Itpr*^*DN*^; *UAS GRAB*_*DA*_ N = 9 brains, 17 cells *p< 0.05, Mann Whitney U test. (lower panel).

### The IP_3_R helps maintain membrane excitability of *PPL1-γ2α′1* neurons

In addition to the neuromodulatory input of acetylcholine, the *PPL1-γ2α′1* DANs are likely to receive direct excitatory inputs. Therefore next we investigated if loss of IP_3_/Ca^2+^ signals alter the essential properties of neuronal excitability of *MB296B* neurons. Stimulation by activation of an optogenetic tool, *CsChrimson* (Klapoetke et al., 2014) was followed by measuring changes in fluorescence of the Ca^2+^ sensor GCaMP in ex-vivo brain preparation. In neurons expressing IP_3_R^DN^ and GCaMP optogenetic stimulation of CsChrimsom did not evince a significant Ca^2+^ response as compared with the control genotype expressing RFP and GCaMP (Figure 6A, B). Similar results were obtained by KCl-evoked depolarisation in the presence of 2 μM Tetrodotoxin (TTX) (Figure 6-figure supplement 1A, B). TTX was added so as to prevent excitation by synaptic inputs from other neurons upon KCl addition. These data suggest that neurons with IP_3_R^DN^ fail to respond to a depolarising stimulus. This idea was tested directly by measuring the change in membrane potential in response to depolarisation, with a genetically encoded fluorescent voltage indicator, *Arclight* (Cao et al., 2013). There was an instant decline in Arclight fluorescence in control cells, while cells with IP_3_R^DN^ showed almost no change in fluorescence after KCl mediated depolarisation (Figure 6C, D). Taken together, these observations confirm that signaling through the IP_3_R is also required for maintaining excitability of central neuromodulatory dopaminergic neurons.

**Figure 6:**
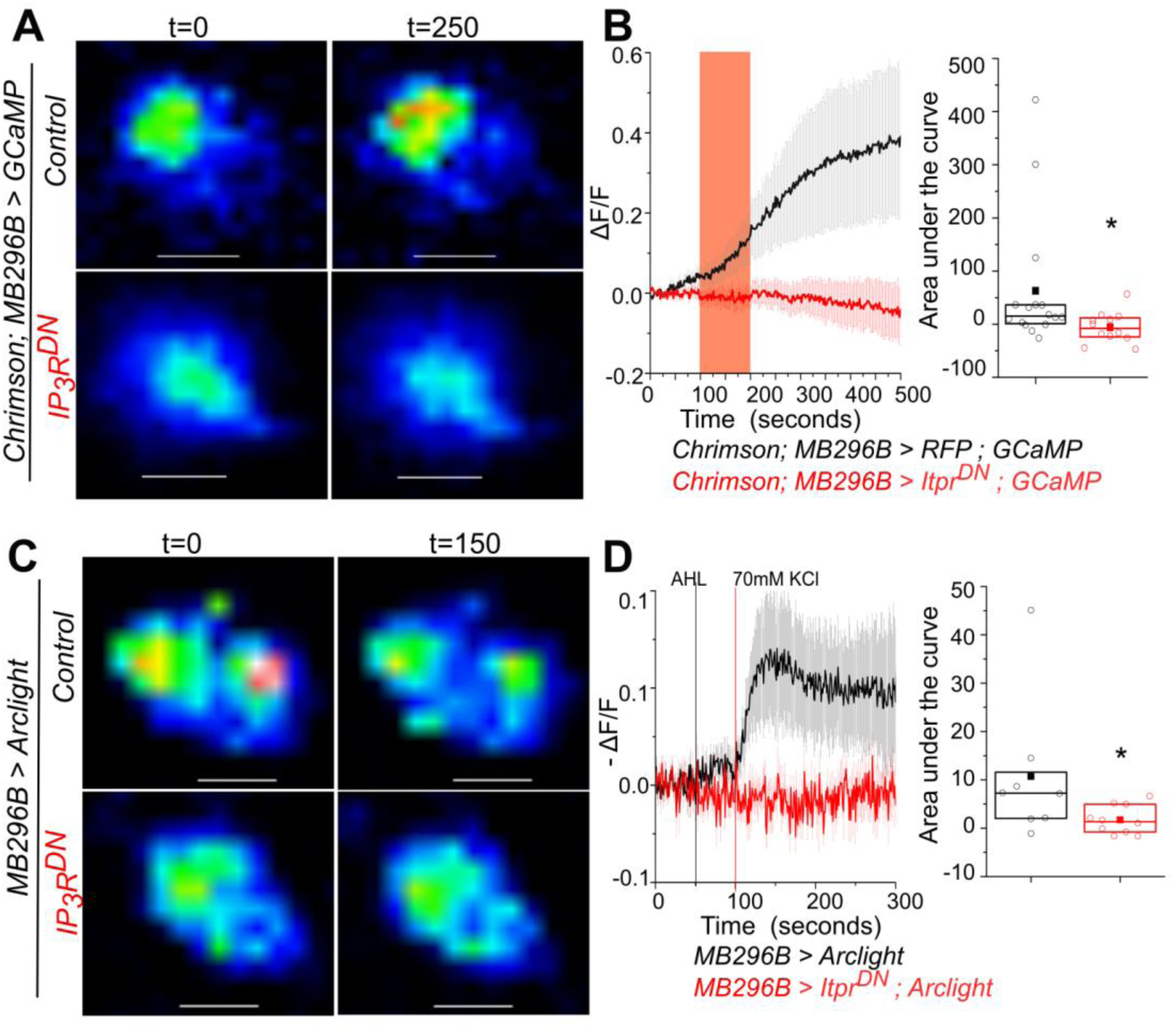
Optimal excitability of PPL1-γ2α′1 dopaminergic neurons requires the IP_3_R. A) Optogenetic activation of PPL1-γ2α′1 DANs, with the red light activated Channelrhodopsin variant Chrimson, is attenuated by expression of IP_3_R^DN^. Representative images of *MB296BGAL4* marked DANs of the indicated genotypes are shown with changes in GCaMP6m fluorescence before (t=0) and after t=250s a 100s pulse of red light (red bar). Images were acquired at 2s per frame. Warmer colors denote increase in [Ca^2+^]. Scale bar = 5 μm (Top). B) Average traces (± SEM) of normalized changes in GCaMP6m fluorescence (ΔF/F) in *MB296BGAL4* marked DANs after activation by Chrimson (left); Quantification of area under the curve from the point of stimulation at 100s up to 400 s. Boxplots and symbols are as described in Figure 2A. *MB296B GAL4 > UAS RFP; UAS GCaMP* N = 7 brains 16 cells; *MB296B GAL4 > UAS Itpr*^*DN*^; *UAS GCaMP* N = 8 brains, 12 cells *p< 0.05, Mann whitney U test. (right). **C)** Changes in membrane potential upon addition of KCl, visualized by expression of the voltage sensor Arclight. Representative images of Arclight responses in *MB296BGAL4* marked DANs are shown from the indicated genotypes and time points. Images were acquired at 1s per frame; Scale bar = 5 μm (Top). **D)** Average traces (± SEM) of normalized changes in Arclight fluorescence (ΔF/F) in *MB296BGAL4* marked DANs after addition of KCl (left). Images were acquired at 1s per frame; Quantification of area under the curve from the point of stimulation at 100s up to 200 s. Boxplots and symbols are as described in Figure 2A. *MB296B GAL4 > UAS Arclight* N = 7 brains 8 cells; *MB296B GAL4 > UAS Itpr*^*DN*^; *UAS Arclight* N = 8 brains, 10 cells *p< 0.05, Mann Whitney U test. (right).

**Figure 6 - figure supplement 1:**
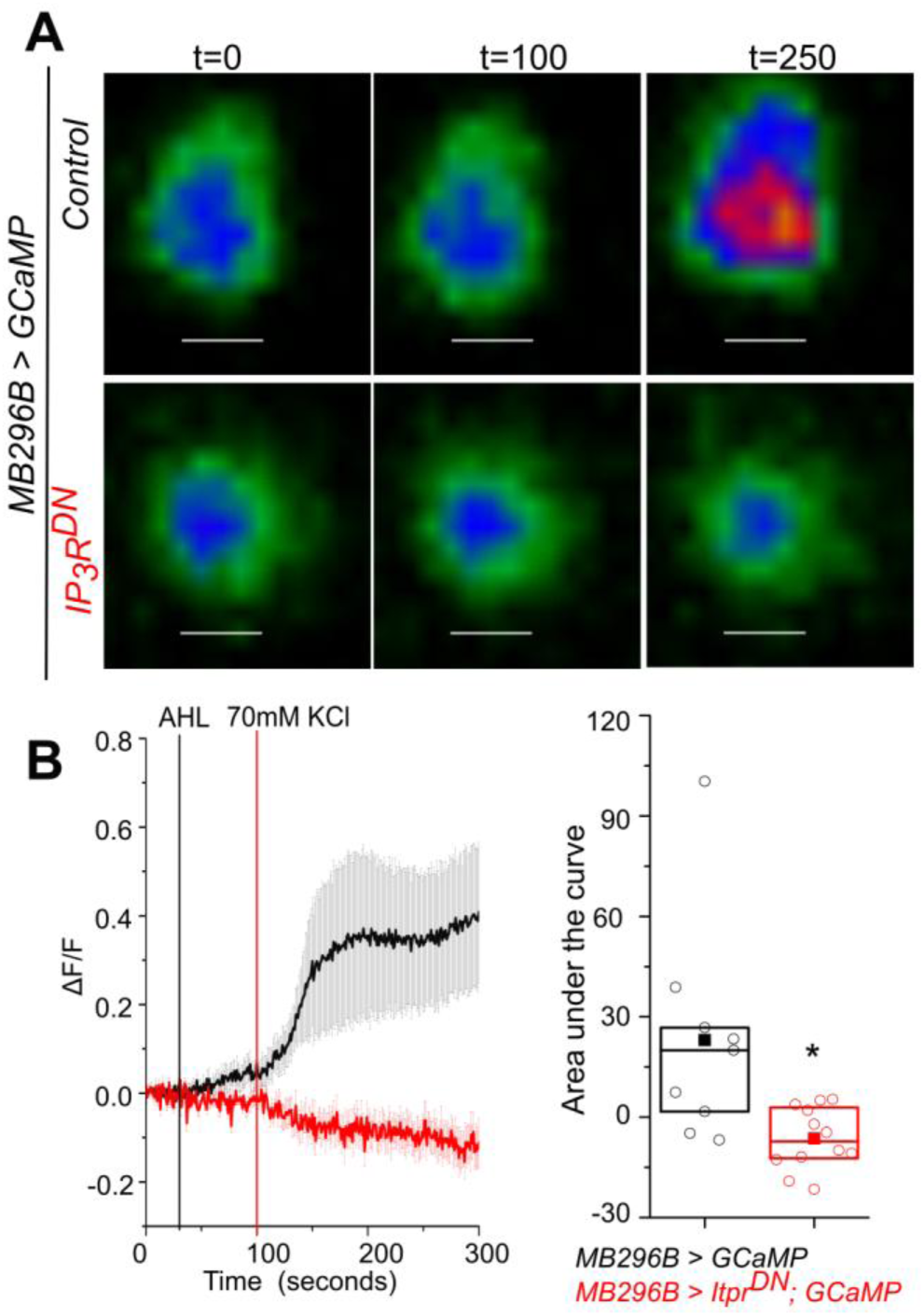
Perturbation of IP_3_R signaling in PPL1-γ2α′1 cells reduced KCl evoked calcium response: A) KCl evoked activation of PPL1-γ2α′1 DANs is attenuated by expression of IP_3_R^DN^. Representative images of *MB296BGAL4* marked DANs of the indicated genotypes are shown with changes in GCaMP6m fluorescence before (t=0, t=100) and after addition of KCl at t=250s. Images were acquired at 1s per frame. Warmer colors denote increase in [Ca^2+^]. Scale bar = 5 μm (Top).) B) Average traces (± SEM) of normalized changes in GCaMP6m fluorescence (ΔF/F) in *MB296BGAL4* marked DANs after activation by KCl (left); Quantification of area under the curve from the point of stimulation at 100s up to 200 s. Boxplots and symbols are as described in Figure 2A. *MB296B GAL4 > UAS GCaMP* N = 5 brains, 9 cells; *MB296B GAL4 > UAS Itpr*^*DN*^; *UAS GCaMP* N = 6 brains, 12 cells *p< 0.05, Mann whitney U test (right).

## Discussion

An inducible IP_3_R^DN^ construct developed and used in this study allowed us to perform stage and cell specific attenuation of IP_3_R mediated calcium signaling *in vivo*. Consistent with well characterised phenotypes of IP_3_R mutants (Banerjee et al., 2004), neuronal expression of IP_3_R^DN^ affected flight. Spatiotemporal studies identified a requirement for the IP_3_R in a small subset of central dopaminergic neurons for maintenance of adult flight bouts as well as in the food-seeking behaviour of hungry flies. Inhibition of synaptic release in the identified dopaminergic subset also reduced the duration of flight bouts. Dopamine release at synapses in the MB γ2α′1 lobe was significantly attenuated in adults expressing the IP_3_R^DN^ (carried over from late pupae). These animals in addition exhibit reduced membrane excitability. Intracellular Ca^2+^ signaling through the IP_3_R is thus required to ensure optimal neuronal excitability and synaptic function in specific central dopaminergic neurons that appear to drive the motivation for both longer flight bouts and the search for food in a hungry fly (Figure 7).

**Figure 7:**
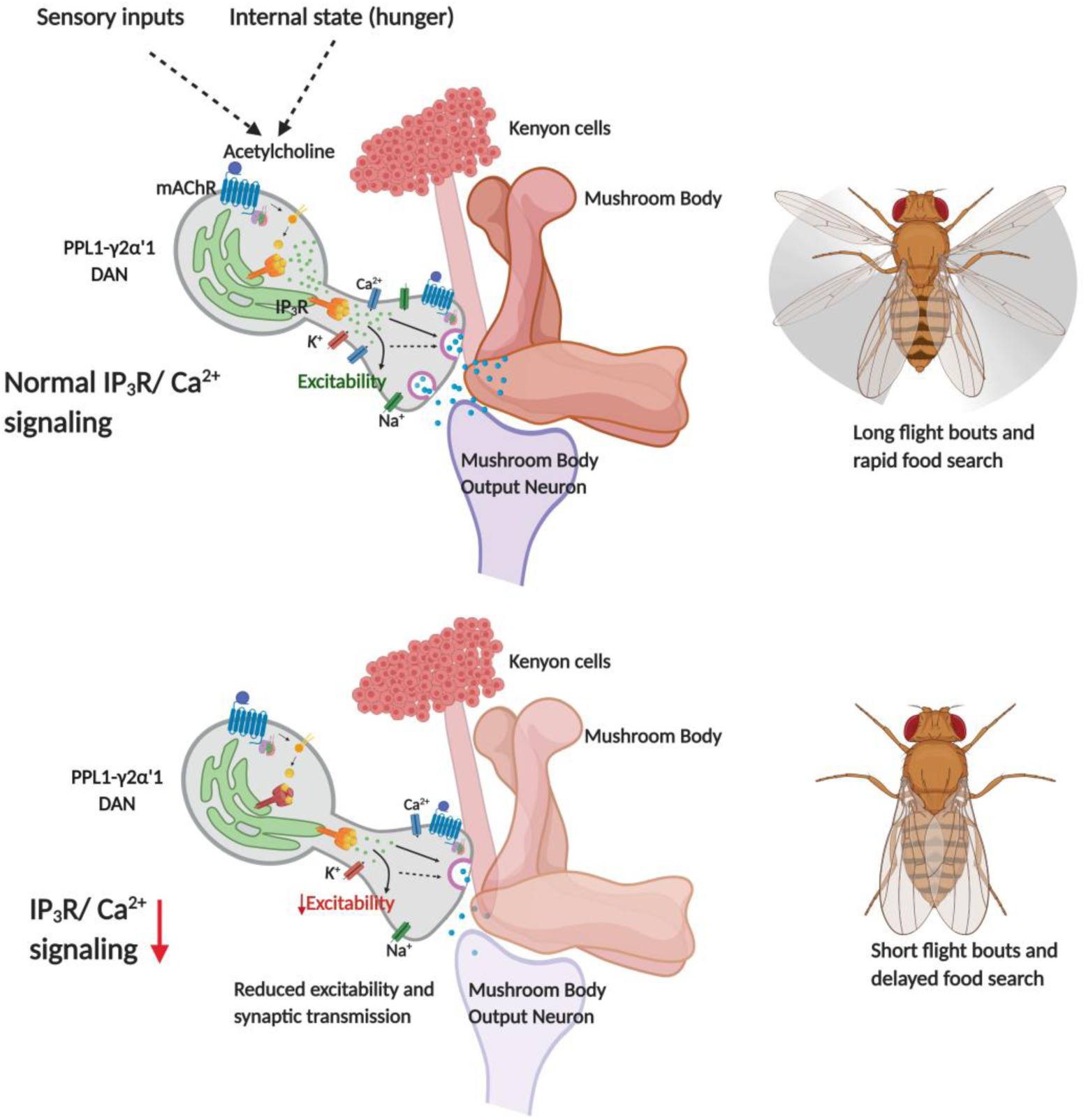
Schematic showing neuronal properties regulated by IP_3_/Ca^2+^ signals in central dopaminergic neurons for flight and food search behaviour in *Drosophila melanogaster*

### The IP_3_R and synaptic release in pupae and adults

Temporal expression of IP_3_R^DN^ demonstrated a requirement during circuit maturation in late pupae but not in adults. However, these data do not rule out an acute function for the IP_3_R in adult dopaminergic neurons because functional wild-type tetramers of the IP_3_R assembled during pupal development very likely mask the effect of adult specific expression of IP_3_R^DN^. Taken together with the short flight bouts observed upon expression of *Shibire*^*ts*^ either in late pupae or in adults, our data support a model where mAChR stimulated ER-Ca^2+^ signals through the IP_3_R regulate synaptic release of dopamine from PPL1-γ2α′1 DANs during circuit maturation and during adult flight. Both direct and indirect effects of ER-Ca^2+^ on synaptic vesicle release have been observed in vertebrates (Gomez et al., 2020; Rossi et al., 2008; Sharma & Vijayaraghavan, 2003) and *Drosophila* (James et al., 2019; Klose et al., 2010; Richhariya et al., 2018; Shakiryanova et al., 2011). Further studies are required to identify the molecular mechanism by which ER-Ca^2+^ signals regulate dopamine release in PPL1-γ2α′1 DANs.

### Intracellular Ca^2+^ signaling and neuronal excitability

Genetic manipulations that target intracellular calcium signaling are known to affect the intrinsic excitability of neurons. For example Purkinje neurons in mice with cell-specific knockout of the ER-Ca^2+^ sensor Stim1, that functions downstream of mGluR1, exhibit a decreased frequency of firing (Ryu et al., 2017). In *Drosophila* FMRFa Receptor mediated calcium signaling modulates the excitability of PPL1 dopaminergic neurons through CAMKII (Ravi et al., 2018). Reduced excitability of dopaminergic neurons expressing the IP_3_R^DN^ transgene (Figure 6 and Figure 6-figure supplement 1) further supports a role for intracellular calcium signals in setting the threshold of membrane excitability in response to neuromodulatory signals, such as acetylcholine (this study), the FMRFa neuropeptide (Ravi et al., 2018) and glutamate (Ryu et al., 2017).

The cellular mechanism(s) by which IP_3_/Ca^2+^ signals regulate neuronal excitability probably vary among different classes of neurons. In addition to the direct activation of Ca^2+^ dependant enzymes such as CamKII, previous reports have shown that knock-down of the IP_3_R in *Drosophila* larval neurons alters their expression profile and specifically affects the expression of several membrane localised ion-channels (Jayakumar et al., 2018). Reduced translation of proteins due to IP_3_R knockdown has also been demonstrated in peptidergic neurons (Megha & Hasan, 2017). Changes in membrane excitability could thus derive from direct regulation of ion channels by Ca^2+^ and Ca^2+^ dependant enzymes as well as an altered density of specific ion channels. Moreover, Ca^2+^ release through the IP_3_R regulates mitochondrial Ca^2+^ entry (Figure 1E). Expression of IP_3_R^DN^ might thus impact neuronal firing and synaptic release by changes in cellular bioenergetics (Cárdenas et al., 2010; Chouhan et al., 2012).

### PPL1-γ2α′1 DANs, flight and the search for food

A role for the PPL1 cluster of dopaminergic neurons in modulating flight and longer flight bouts has been reported earlier (Pathak et al., 2015; Ravi et al., 2018). However, amongst the PPL1 cluster this is the first report identifying the two PPL1-γ2α′1 DANs as required for maintenance of flight. The PPL1-γ2α′1 DANs and their downstream Mushroom body output neuron (MBON-γ2α′1) also encode the state of hunger (Tsao et al., 2018), formation and consolidation of appetitive memory (Berry et al., 2018; Felsenberg et al., 2017; Yamazaki et al., 2018) and sleep (Aso et al., 2014; Sitaraman et al., 2015). These studies identified the importance of modulated dopamine release as a motivational cue wherein dopamine release from PPL1-γ2α′1 DANs increased in starved flies (Tsao et al., 2018) and inhibited the activity of a cholinergic output neuron from the Mushroom Body (MBON-γ2α′1; Tsao et al., 2018). The MBON-γ2α′1 projects to another set of central DANs, the PAM neurons (Felsenberg et al 2017), identified as flight promoting in a previous study (Manjila et al., 2019). This leads to the hypothesis that inputs to the PPL1-γ2α′1 DANs through mAChR driven IP_3_/Ca^2+^ signals modulate dopamine release at the PPL1-γ2α′1> MBON-γ2α′1 synapse, and the extent of dopamine release changes the output strength of MBON-γ2α′1. Altered MBON-γ2α′1 outputs might then regulate flight through the PAM-DANs and their downstream circuits. Prolonged flight bouts are energy intensive and very likely require multisensory integration for their continuation. Modulated activity in the PPL1-γ2α′1 DANs forms an important component of this integration and can lead to inhibition of MBON-γ2α′1 so as to maintain wakefulness (Aso et al., 2014; Sitaraman et al., 2015) as well as increase the search for food when hungry (Tsao et al., 2018). The integration of external cues that stimulate flight with the internal states of hunger and wakefulness very likely serve important functions of survival in the wild.

## Materials and methods

### Fly stocks

*Drosophila* strains used in this study were reared on cornmeal media, supplemented with yeast. Flies were maintained at 25°C, unless otherwise mentioned under 12:12 light: dark cycle. WT strain of *Drosophila* used was *Canton S*.

Following fly lines were kindly provided by various fly community members across the globe as mentioned: *THGAL4* (Friggi-Grelin et al., 2003) by Serge Birman (CNRS, ESPCI Paris Tech, France), *THD’* GAL4 (Liu et al., 2012) from Mark N Wu (Johns Hopkins University, Baltimore), *UAS GRAB*_*DA*_ (Sun et al., 2018) from Youlong Li (Peking University School of Life Sciences, Beijing, China), *UAS Shibire*^*ts*^ (Vicario et al., 2001) from Toshihiro Kitamoto (University of Iowa, Carver College of Medicine, Iowa). *vGlut*^*VGN6341*^*GAL4* (Syed et al., 2016) from K. Vijayraghavan (NCBS, India), *UAS mitoGCaMP* (Lutas et al., 2012) from Fumiko Kawasaki (Pennsylvania State University, Pennsylvania). A strain with two copies of *TubGAL80*^*ts*^ on the third chromosome was generated by Albert Chiang, NCBS, Bangalore, India, and has been used for all *Gal80*^*ts*^ experiments. Single-point mutants of the *itpr* gene (*Itpr*^*ka1091*^, *itpr*^*ug3*^) were characterized as described previously (Joshi et al., 2004)

The fly lines *nSybGAL4* (BL51635), *UAS GCaMP6m* (BL42748), *UAS Arclight* (BL51057), *UASmCD8GFP* (BL5130), *UAS Dicer* (BL24648), *UAS Chrimson* (BL55136), *UAS syt*.*eGFP* (BL6926), *MB058BGAL4* (BL68278), *MB296BGAL4* (BL63308), *MB304BGAL4* (BL68367), *MB320CGAL4* (BL68253), *MB630BGAL4* (BL68334), *MB438BGAL4* (BL68326), *MB504BGAL4* (BL68329) were obtained from Bloomington Drosophila Stock Centre (BDSC). The *UAS itprRNAi* (1063-R2) strain was from National Institute of Genetics (NIG), *UAS mAChR RNAi* was from VDRC (101407)

### Single flight assay

Flight assays were performed according to Manjila & Hasan, 2018. Briefly, 3-5 day old flies of either sex were tested in batches of 8-10 flies. They were anaesthesized on ice for 2-3 minutes and then tethered between their head and thorax using a thin metal wire and nail polish. Once recovered, mouth blown air puff was given as a stimulus to initiate flight and flight time was recorded for each fly till 15 minutes. For all control genotypes, *GAL4* or *UAS* strains were crossed to wild type strain, *Canton S*. Flight time data is represented in the form of boxplots using Origin software (OriginLab, Northampton, MA). Each box represents 25th to 75th percentile, each open circle represents flight duration of a single fly, solid squares represent the mean and the horizontal line in each box represents the median.

For Gal80^ts^ experiments, larvae, pupae or adults were maintained at 18°C and transferred to 29°C only at the stage when the *UAS* transgene needed to be expressed. Flight assay was done at 25°C. For adult specific expression, flies were grown at 18°C and transferred to 29°C immediately after eclosion for 2-3 days. For experiments involving *Shibire*^*ts*^, larvae, pupae or adults were maintained at 22°C and transferred to 29°C only at the stage when the *Shibire*^*ts*^ needs to be activated. Flight assay was done at 25°C except for adult specific activation in which flies were shifted to 29°C ten minutes before flight assay and then maintained at 29°C during the experiment.

For optogenetic experiments, flies were transferred to media containing 200 mM all-trans-retinal (ATR) and reared in dark for 2-3 days before imaging experiments.

### Ex vivo live Imaging

Adult brains were dissected in Adult hemolymph-like (AHL) saline (108 mM NaCl, 5 mM KCl, 2 mM CaCl2, 8.2 mM MgCl2, 4 mM NaHCO3, 1 mM NaH2PO4, 5 Mm trehalose, 10 mM sucrose, 5 mM Tris, pH 7.5) while larval brains were dissected in HL3 (70 mm NaCl, 5 mm KCl, 20 mm MgCl2, 10 mm NaHCO3, 5 mm trehalose, 115 mm sucrose, 5 mm HEPES, 1.5 mm Ca2+, pH 7.2). The dissected brain was mounted on culture dish with anterior side up for recording from cell while posterior side up for imaging mushroom body. They were then embeded in 6µl of 1% low-melt agarose (Invitrogen), and bathed in AHL. Images were taken as a time series on an XY plane using a 20x objective on an Olympus FV3000 inverted confocal microscope (Olympus Corp.). Acquisition time is different for different experiments and is described in the figure legends. GCaMP6m, Arclight and GRAB_DA_ signals were captured using the 488 nm excitation laser line while 633nm laser was used for optogenetic stimulation of Chrimson.

Raw fluorescence data were extracted from the marked ROIs using a time series analyzer plugin in Fiji (Balaji, https://imagej.nih.gov/ij/plugins/time-series.html). ΔF/F was calculated using the following formula for each time point (t): ΔF/F = (F_t_-F_0_)/F_0_, where F_0_ is the average basal fluorescence of the first 20 frames.

To quantify response to stimuli we calculated area under the curve(AUC). Area under the curve was calculated from the point of stimulation till mean peak response was reached using Microsoft Excel (Microsoft). Time frame for calculating AUC is mentioned in figure legends. AUC is represented as boxplots using Origin software (OriginLab, Northampton, MA). Each box represents 25th to 75th percentile, each open circle represents flight duration of a single fly, solid squares represent the mean and the horizontal line in each box represents the median.

### Generation of IP_3_R^DN^

Five *itpr* residues in *Drosophila itpr* cDNA (Sinha & Hasan, 1999) were mutated using site directed mutagenesis kit (Agilent). The oligonucleotide CAGAGATCGGCAG**C**AATTGCTGC**AG**GAACAGTACATCC was used to change K530/R533 to Q while GTACCACGTCTTTCTGC**AG**ACCACCGGACGCACCAG was used to change R272 to Q. All mutations were confirmed using Sanger’s sequencing. Mutated *itpr* cDNA was subcloned in *UAS attB* vector (Bischof et al., 2007). *UAS Itpr*^*DN*^ plasmid was then microinjected in fly embryos at the NCBS fly facility to obtain stable fly strains using standard protocols of fly embryo injection.

### Food seeking assay

Food seeking assay was performed according to Tsao et al., 2018. Briefly, 18-20 hrs starved males (4 days old) of specified genotypes were introduced in petridish (dimensions) with drop of yeast solution in the centre. The yeast solution was prepared by mixing 0.2 gram of yeast with 1 gram of sugar in 5 ml distilled water and incubated in a 28°C shaking incubator (170 rpm) for 16 hr. Starved males were then allowed to search for food and it was considered having found food if it rested for 3 sec or longer on food drop. Food-seeking index was calculated as: [Total assay time (600 s) - the time taken to locate food (sec)]/Total assay time (600 s).

### Immunohistochemistry

Immunohistochemistry was performed on dissected adult brains as described in Pathak et al., 2015. Briefly, brains were dissected in 1x PBS, followed by fixation in 4% paraformaldehyde for 30 minutes at room temperature and then 3-4 washings with 0.2% phosphate buffer, pH 7.2 containing 0.2% Triton-X 100 (PTX). They were then blocked in 0.2% PTX containing 5% normal goat serum for four hours at 4°C and incubated overnight with primary antibodies. Next day, they were washed three to four times with 0.2% PTX at room temperature, and then incubated with the respective fluorescent secondary antibodies for 2 hr at room temperature. The primary antibodies used were: rabbit anti GFP (1:10,000, A6455, Life Technologies), mouse anti-bruchpilot (anti-brp) antibody (1:50; kindly provided by Eric Buchner, Univerity of Wuerzburg, Germany). Fluorescent secondary antibodies used at were anti-mouse Alexa Fluor 568 (1:400, #A11004, Life Technologies) and anti-rabbit Alexa Fluor 488 (1:400, #A11008, Life Technologies). Confocal images were obtained on the Olympus Confocal FV3000 microscope (Olympus Corp.) with a 20x or with a 40x objective. Images were visualized using Fiji. For Syt.eGFP fluorescence, mean intensity fluorescence was obtained for syt.eGFP and mRFP by averaging intensity values between anterior and posterior limits of the structure.

### Western Blots

Between 5 to 10 pupal and adult brains or adult heads of appropriate genotypes were dissected in cold PBS and were homogenized in 30 μl of homogenizing buffer (25 mM HEPES, pH 7.4, 150 mM NaCl, 5% glycerol, 1 mM DTT, 1% Triton X-100, and 1 mM PMSF). 15 μl of the homogenate was run on a 5% SDS-polyacrylamide gel. The protein was transferred to a nitrocellulose membrane by standard protocols. Membrane was then blocked with 5% skim milk followed by incubation with primary antibody at 4°C overnight. The affinity-purified anti-IP_3_R rabbit polyclonal antibody (Agrawal et al., 2009) (IB-9075) was used at a dilution of 1:300 and mouse anti-spectrin antibody (1:50; 3A9, DSHB) was used as a loading control for IP_3_R. Secondary antibodies conjugated with horseradish peroxidase were used at dilution of 1:3000 (anti-mouse HRP; 7076S, Cell Signaling Technology) and 1:5000 (anti-rabbit HRP; 32260, Thermo Scientific). The protein was then detected on the blot by a chemiluminiscent detection solution (Advansta). First spectrin antibody was used to detect protein. Blot was then washed with 3% glacial acetic acid and reprobed with IP_3_R antibody.

### Statistical tests

For flight data, two-tailed Student’s t test was used to compare two samples while one-way ANOVA followed by post hoc Tukey’s test was used if more than two conditions were involved. All statistical tests were performed using Origin 8.0 software (Micro Cal). Statistical tests and p-values are mentioned in each figure legend. For food seeking assay as well as imaging data, Two-tailed Mann-Whitney U test was performed to statistically validate the significance.

Model in Figure 7 was created using Biorender (BioRender.com)

## ACKNOWLEDGMENTS

This study was supported by grants from DBT-SERB and NCBS-TIFR to G.H. A.S. was supported by a fellowship from the National Centre for Biological Sciences, TIFR. We thank the Fly Facility, Sequencing facility and Central Imaging and Flow Cytometry Facility at NCBS.

